# Early adversity promotes adolescent avoidance behavior by enhancing prefrontal–amygdala communication through CRH⁺ glutamatergic neurons

**DOI:** 10.64898/2025.12.23.696229

**Authors:** Caitlin M. Goodpaster, Zachary Zeidler, Michael W. Gongwer, Charles J. Bland, Meelan Shari, Cassandra B. Klune, Nico S. Jones, Makayla Ramirez, Maryam-Batul Alturki, Rio Hundley, Jolie Utter, Laura A. DeNardo

## Abstract

Early life adversity (ELA) has lasting impacts on emotion circuit function^1^. Exposure to early maltreatment, neglect, or household instability is associated with enduring alterations in threat evaluation and response, hallmarks of mood and anxiety disorders^2–4^. These alterations often emerge during adolescence and are characterized by excessive avoidance that limits engagement in rewarding experiences^5,6^. However, the neurodevelopmental mechanisms linking ELA to increased threat responses remain unclear. We identified a stress-sensitive medial prefrontal cortex (mPFC)-basolateral amygdala pathway that co-expresses corticotropin-releasing hormone (CRH) and glutamate. We find that ELA strengthens this pathway and increases threat avoidance behavior during adolescence. Optogenetic inhibition and developmental CRH knockdown in this pathway both rescue ELA-induced elevations in adolescent avoidance, whereas broadly reducing mPFC CRH expression in typically reared mice mimics ELA phenotypes. The same manipulations produced no behavioral effects in adults. Together, our findings indicate that cell type-specific CRH signaling regulates developmental circuit plasticity, biasing emotional circuit function and potentially contributing to psychiatric vulnerability.

## INTRODUCTION

Early life adversity (ELA), including neglect, abuse, maltreatment, or an unstable home environment, affects nearly 60% of people worldwide^7–10^. Although ELA may be transient, it induces lasting changes in the development and function of the brain’s emotion systems, including how individuals evaluate and respond to environmental threats^1^. As such, ELA is a major risk factor for psychiatric conditions, including anxiety and depression^2–4^. These disorders often emerge during adolescence, a critical stage of brain development, when exploration and experience shape late-developing cognitive and emotional systems through activity-dependent plasticity^5,6^. Mood and anxiety disorders are characterized by heightened threat avoidance that can disrupt neural circuit maturation and interfere with goal-directed behaviors^11,12^. Despite this, most mechanistic studies of ELA focus on adult outcomes, and the developmental processes linking ELA to later behavioral changes remain poorly understood^1,11,13^. This information can ultimately help us identify, prevent and treat psychiatric symptoms in at-risk populations.

The limbic system, which processes emotional stimuli and regulates learning and decision making^14,15^, is particularly sensitive to adversity due to its high expression of stress hormone receptors^16^. It is also dysregulated in psychiatric disorders^11,17–19^. Human neuroimaging studies reveal that ELA alters limbic system activity and functional connectivity, especially between the medial prefrontal cortex (mPFC) and basolateral amygdala (BLA)^11,20–22^. Although some of these alterations correlate with maladaptive threat responses^20,23,24^, the absence of causal evidence leaves key gaps in understanding how ELA disrupts the development of relevant brain circuits to cause exaggerated threat avoidance. As behavior arises from the coordinated activity of specific neural circuits in the brain, uncovering the cell type- and circuit-specific changes downstream of ELA will ultimately reveal the mechanisms by which ELA impacts cognitive and emotional behaviors.

Stress-sensitive molecules such as corticotropin-release hormone (CRH) are enriched in specific limbic cell populations and may link ELA to persistent behavioral changes^25–27^. Most studies have focused on CRH in the hypothalamus, where it regulates the body’s stress response^26,28^. However, CRH is also highly expressed in key emotion-regulatory centers, including the mPFC, hippocampus, central amygdala (CeA), and bed nucleus of the stria terminalis (BNST)^29^. ELA alters both CRH expression in these regions and the function of CRH-expressing neurons. CRH is primarily expressed in Gamma-Aminobutyric Acid (GABAergic) cells^28,30^ and signals through its cognate receptors—corticotropic releasing factor receptors 1 and 2—to promote plasticity in target neurons^31^. Reducing CRH activity, either through gene knockdown or pharmacological receptor inhibition, in the CeA, BNST, and hippocampus respectively rescues ELA-induced anhedonia, anxiety-like behaviors, and memory deficits^32–34^. In addition, inhibiting CRH-expressing CeA cells prevents ELA-induced increases in startle responses and maladaptive reward behaviors in adulthood^35,36^.

Together, these findings highlight CRH as both an effector that drives stress-induced neuroplasticity and a marker of stress-sensitive pathways that contribute to the enduring consequences of ELA. However, we have a limited understanding of how ELA-induced alterations in the CRH system influence brain development— particularly within the circuits that support complex threat responses, including avoidance behaviors that are central to anxiety disorders. Most studies of the effects of ELA on the CRH system focused exclusively on adults^32,33,36,37^. In addition, many of these studies that broadly manipulated CRH or receptor expression produced conflicting findings, highlighting the need for region- and cell type-resolved studies of CRH circuit function^38^. Revealing how ELA uniquely impacts specific CRH circuits in the developing brain can provide a foundation for developing more effective interventions for psychiatric illnesses that are informed by age-specific biology.

Using a mouse model of ELA, we observed elevated threat avoidance behavior in adolescence that was correlated with heightened BLA activity. Brain-wide mapping of CRH revealed that ELA induces a selective decrease in the number of CRH+ neurons in the ventromedial (vm) PFC in adolescents. In line with this, vmPFC-wide CRH knockdown earlier in development reproduces ELA phenotypes in typically reared adolescent mice. Using viral circuit mapping and patch clamp recordings, we identified a glutamatergic, CRH-expressing neuronal population that projects from the vmPFC to the BLA and is potentiated following ELA. RNAscope revealed that while ELA reduces the number of non-glutamatergic vmPFC CRH+ cells, it drives higher CRH expression in the glutamatergic CRH+ population. In line with this, optogenetic inhibition of the glutamatergic vmPFC^CRH+^→BLA pathway rescues ELA-induced elevations in threat avoidance in adolescents. Similarly, CRH knockdown selectively in excitatory vmPFC cells leads to reduced avoidance and BLA activity in adolescent ELA mice. In adulthood these manipulations had no effect, highlighting the developmentally restricted role of this pathway in driving heightened avoidance following adversity. Together, these findings suggest that vmPFC CRH functions as an early postnatal effector following ELA, acting in a cell-type specific manner and inducing plasticity in an excitatory, CRH-expressing vmPFC–BLA pathway. By elucidating how ELA transforms activity in a previously undescribed CRH-expressing excitatory pathway, this work provides mechanistic insight into how ELA alters adolescent behavior and may inform strategies to mitigate risk for mental illness.

## RESULTS

### ELA augments adolescent threat avoidance behavior and underlying BLA activity

Limbic areas, including the mPFC and BLA, regulate threat avoidance behaviors in adults^39,40^. These circuits undergo a marked rearrangement during adolescence, contributing to age-dependent changes in threat avoidance behavior^40^. Compared with adults, adolescent mice typically display lower levels of threat avoidance behavior, instead promoting exploration^40,41^. ELA amplifies threat responses in both rodents and humans, an effect that correlates with increases in BLA activity^20,42,43^. However, it remains unknown how ELA alters learned responses to avoidable threats and what cellular and molecular mechanisms underlie these changes. We hypothesized that ELA-induced alterations in limbic circuits would be pronounced during adolescence, driving behavioral changes during this developmental window when psychiatric symptoms of anxiety and depression, including heightened threat responses, typically emerge.

To test this hypothesis, we used the limited bedding and nesting (LBN) model of ELA^35,44,45^. LBN mimics a low resource environment by reducing the dam’s nesting material and separating mice from the bedding material with a wire mesh floor (Fig. 1a). This provokes fragmented and unpredictable maternal care during a critical developmental window (postnatal day (P)4—P11)^46^. During this period, LBN dams spent less time nursing and more time alone (Fig. 1b), resulting in an overall reduction in time on the nest (Fig. 1c). LBN pups also weighed less than standard reared (SR) controls at P12, but this difference was no longer evident by weaning (Fig. 1d).

**Figure 1.**
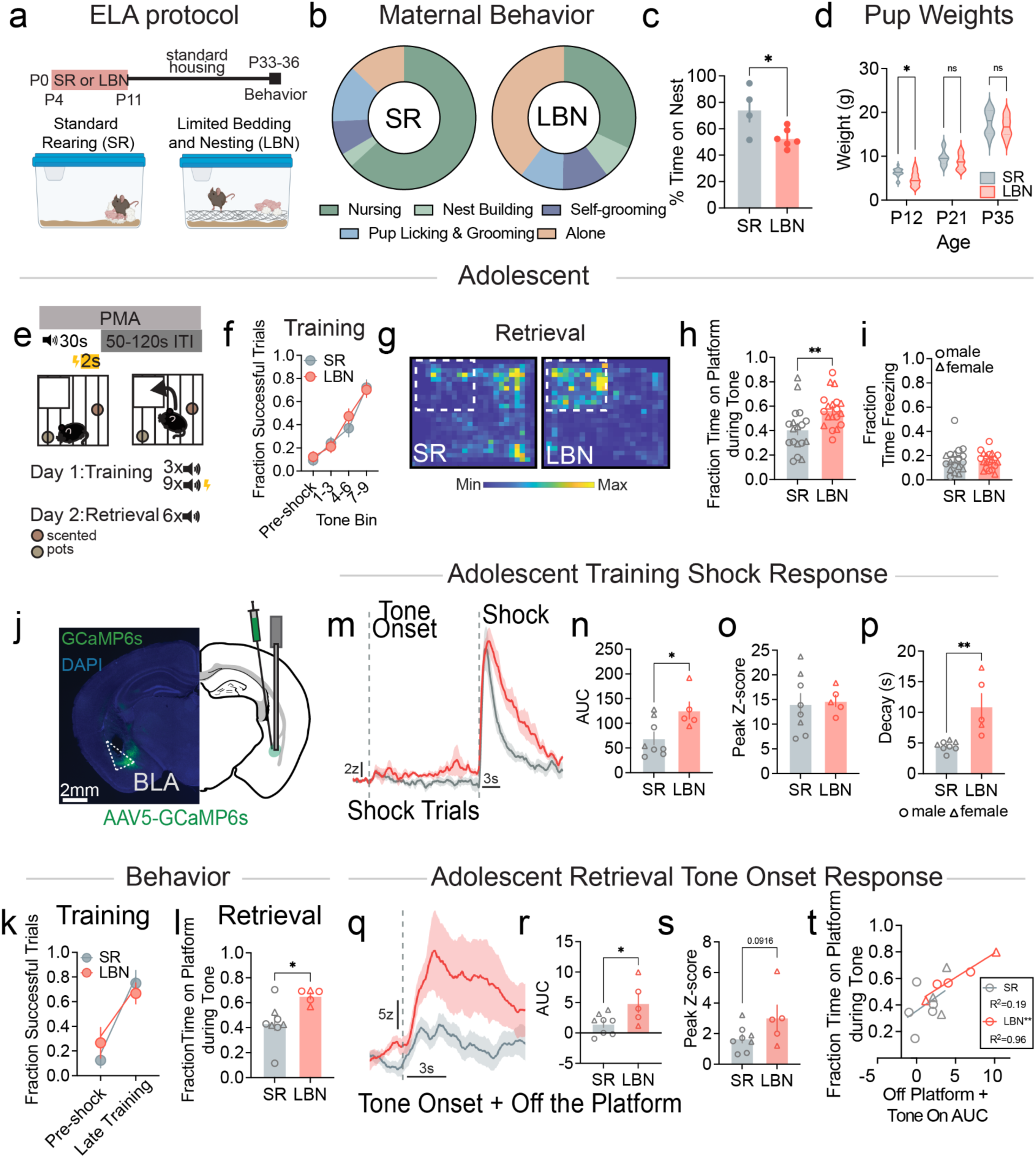
ELA augments adolescent threat avoidance behavior and underlying BLA activity. (**a**) LBN protocol. (**b**) Distribution of maternal behaviors during LBN versus standard rearing (SR) (SR, n=4; LBN n=6 dams) (Chi-squared test ꭓ^2^(4,27), p<0.0001). (**c**) Percent time dams spend on the nest (unpaired t-test F_3,5_=6.51, p=0.028). (**d**) Weight of pups at postnatal day (P)12, P21, P35 (SR, n=16 pups/4 litters, LBN, n=15 pups/6 litters; two-way repeated measured ANOVA with Bonferroni’s post hoc test: age effect F_1.51,43.92_ = 736.4, P<0.0001; rearing effect *F*_1,29_ = 3.53, *P*=0.071; interaction *F*_1.51,43.92_ = 0.1949, *P* = 0.762, violin plots display median and quartiles). (**e**) Platform Mediated Avoidance (PMA) protocol. (**f**) Fraction of successful trials, averaged across bins of 3 tones (SR, n=18; LBN, n=19 mice; two-way repeated measures ANOVA with Sidak’s post hoc test: trial effect *F*_2.29,80.10_ = 40.84, *P*<0.0001; rearing effect *F*_1,35_ = 0.22, *P*=0.645; interaction *F*_2.29,80.10_ = 0.55, *P* = 0.605). (**g**) Representative examples of mouse location depicted as heatmaps during PMA retrieval. (**h**) Averaged fraction of time on the platform during the tone during retrieval (unpaired t-test *F*_17,18_=1.71, *P*=0.006). (**i**) Averaged fraction time spent freezing during the tone during retrieval (unpaired *t*-test *F*_17,18_=2.32 p=0.888). (**j-t**), Fiber photometry recordings in BLA during PMA training and retrieval (SR, n=8; LBN, n=5 mice). (**j**) Schematic and example images of GCaMP6s expression and fiber optic implant in the BLA. (**k**) Fraction of successful trials during PMA training of fiber photometry animals (SR, n=8; LBN, n=5 mice) (2-way ANOVA: trial effect *F*_1,11_=30.09, *P*=0.0002; rearing effect *F*_1,12_=0.09, *P*=0.77; interaction effect *F*_1,11_=1.45, *P*=0.254). (**l**) Fraction of time on platform during PMA retrieval of photometry animals (unpaired *t*-test *F*_7,4_=8.10, *P*=0.016). (**m**) Averaged change in fluorescence (z-score) in BLA activity during the tone periods in which the mouse received a shock during PMA training. (**n**) Quantification of area under the (AUC) during the shock and 13 seconds afterwards (unpaired t-test *F*_4,7_=1.07, *P*=0.024). (**o**) Quantification of peak fluorescence during shock (unpaired t-test *F*_7,4_=5.10, *P*=0.829). (**p**) Quantification of decay rate (seconds) of exponential curve fit to the return to baseline following shock (unpaired t-test *F*_4,7_=33.56, *P*=0.0026). (**q**) Averaged change in fluorescence (z-score) in BLA activity in response to tone onset when the mouse was off the platform. (**r**) Quantification of AUC during the 3 seconds following tone onset (unpaired t-test *F*_4,7_=4.36, *P*=0.037). (**s**) Quantification of peak response during the 3 seconds following tone onset (unpaired t-test *F*_4,7_=5.06, *P*=0.092). (**t**) Correlation between AUC and fraction of time on the platform during the tone (linear regression SR: *F*_1,6_=1.44, *P*=0.276, LBN: *F*_1,3_=66.41, *P*=0.004). All data points represent biological replicates. Data are shown as mean ± s.e.m., and shading reflects between-subjects s.e.m.; \**P*<0.05, \*\**P*<0.01. Triangles denote females, and circles denote males.

To assess changes in threat avoidance behavior during adolescence (P33-37), we used the platform mediated avoidance (PMA) assay^40,47^, where mice learn to move to a platform to avoid a tone-cued shock (Fig. 1e). Odor pots placed out of reach below the shock bars encouraged exploration of the chamber. While LBN mice showed slightly reduced sensitivity to the shock (Extended Data Fig. 1a), this did not impact their ability to learn PMA. Both groups reached similar performance levels by the end of training, as measured by the fraction of successful trials and the time on the platform during the tone (Fig. 1f, Extended Data Fig. 1b). In contrast, during a threat memory retrieval session the following day, LBN mice spent significantly more time on the platform during the tone (Fig. 1g,h), tending to take less time to enter the platform after tone onset (Extended Data Fig. 1c). Importantly, freezing levels did not differ between groups (Fig. 1i), indicating that the increased platform time was not driven by heightened freezing at this location or by a stronger tone-shock association, but instead by differences in threat-guided avoidance behavior.

To determine whether increased PMA was driven by increased anxiety-like behaviors in LBN mice, we used the open field and the elevated plus maze (Extended Data Fig. 1d,e). These assays measure how much time mice spend in exposed parts of an apparatus, mimicking where they are more vulnerable to predation. We did not observe rearing-related behavioral changes in these assays suggesting that ELA induces learning-specific changes in threat responding, rather than more general increases in anxiety-like behaviors. No differences were observed in a novel odor assay, suggesting that ELA does not affect the drive to explore, but increases avoidance specifically when exploration carries potential threat or cost (Extended Data Fig. 1f).

To assess how ELA may alter BLA activity underlying threat avoidance in adolescent mice, we used fiber photometry to measure bulk calcium fluorescence, a proxy for neural activity, during both PMA training and retrieval (Fig. 1j). Following PMA recordings, correct viral expression and fiber placement were confirmed in all animals (Extended Data Fig. 2a). Similar levels of learning were observed in both groups, while LBN mice exhibited significantly increased avoidance during retrieval (Fig. 1k,l). During foot shocks, SR and LBN groups exhibited shock responses of similar amplitude (Fig. 1m). However, in LBN mice, these responses returned to baseline more slowly, resulting in an overall increase in shock-evoked Ca^2+^ activity compared to SR controls (Fig. 1n-p). During retrieval, the tone onset response in the LBN groups was significantly greater than controls, particularly when the animals were off the platform and the threat was imminent (Fig. 1q-s). In LBN but not control mice, avoidance levels were robustly correlated with this tone onset response (Fig. 1t), suggesting that the tone-related BLA activity strongly encodes cued threat avoidance in LBN mice but not in controls.

In separate cohorts of mice, we further investigated the cellular and synaptic changes that may underlie the enhanced BLA activity in LBN mice. We first measured the expression levels of phosphorylated cAMP response element binding protein (pCREB), a transcription factor linked to neuronal excitability^48^. LBN mice had significantly elevated pCREB expression levels, despite no changes in BLA cell number and no differences in PMA learning (Extended Data Fig. 2b-e). We next recorded spontaneous excitatory and inhibitory postsynaptic currents (sEPSCs and sIPSCs, respectively) from BLA neurons in acute brain slices (Extended Data Fig. 2f-h). LBN mice showed an overall trend toward higher sEPSC frequency and exhibited a significantly higher proportion of high amplitude sEPSCs (Extended Data Fig. 2i), suggesting LBN leads to enhanced excitatory synaptic input into BLA neurons. In contrast, we observed no changes in IPSCs (Extended Data Fig. 2j), suggesting that the effects of LBN were specific to excitatory synapses. Together, these results suggest that ELA may heighten BLA activity during adolescent threat avoidance by increasing neuronal excitability and excitatory synaptic input.

### Reduced CRH in the vmPFC is associated with heightened threat avoidance in adolescence

CRH is a central mediator of stress and threat processing^26,28^. Infusion of CRH into the BLA increases activation of excitatory populations and enhances fear memories^49–51^ and stress impacts CRH receptor activity in the BLA^7,52^. However, BLA expresses little CRH itself^29^, suggesting altered CRH release originates from an external source, yet it is unknown how ELA impacts CRH expression in BLA-projecting areas during adolescence.

To systematically examine how ELA affects CRH populations across the adolescent brain, we crossed *CRH-Cre*^53^ mice with the Ai14 reporter line^54^ so that all CRH neurons would be labeled with a red fluorophore (tdTomato). Following LBN or SR, we used whole-mount tissue clearing, immunolabeling, and light sheet microscopy to quantify the number of CRH-expressing (CRH+) cells in every brain region for each rearing group (Fig. 2a). Consistent with the previous literature^29^, we observed dense clusters of CRH+ cells in the CeA, piriform cortex (PIR), BNST and hypothalamus (Fig. 2b). We quantified CRH+ cell density changes in regions known to send the most robust projections to BLA^55^ and observed a selective reduction in CRH^+^ cell density in the vmPFC (Fig. 2c, Extended Data Table 1).

**Figure 2.**
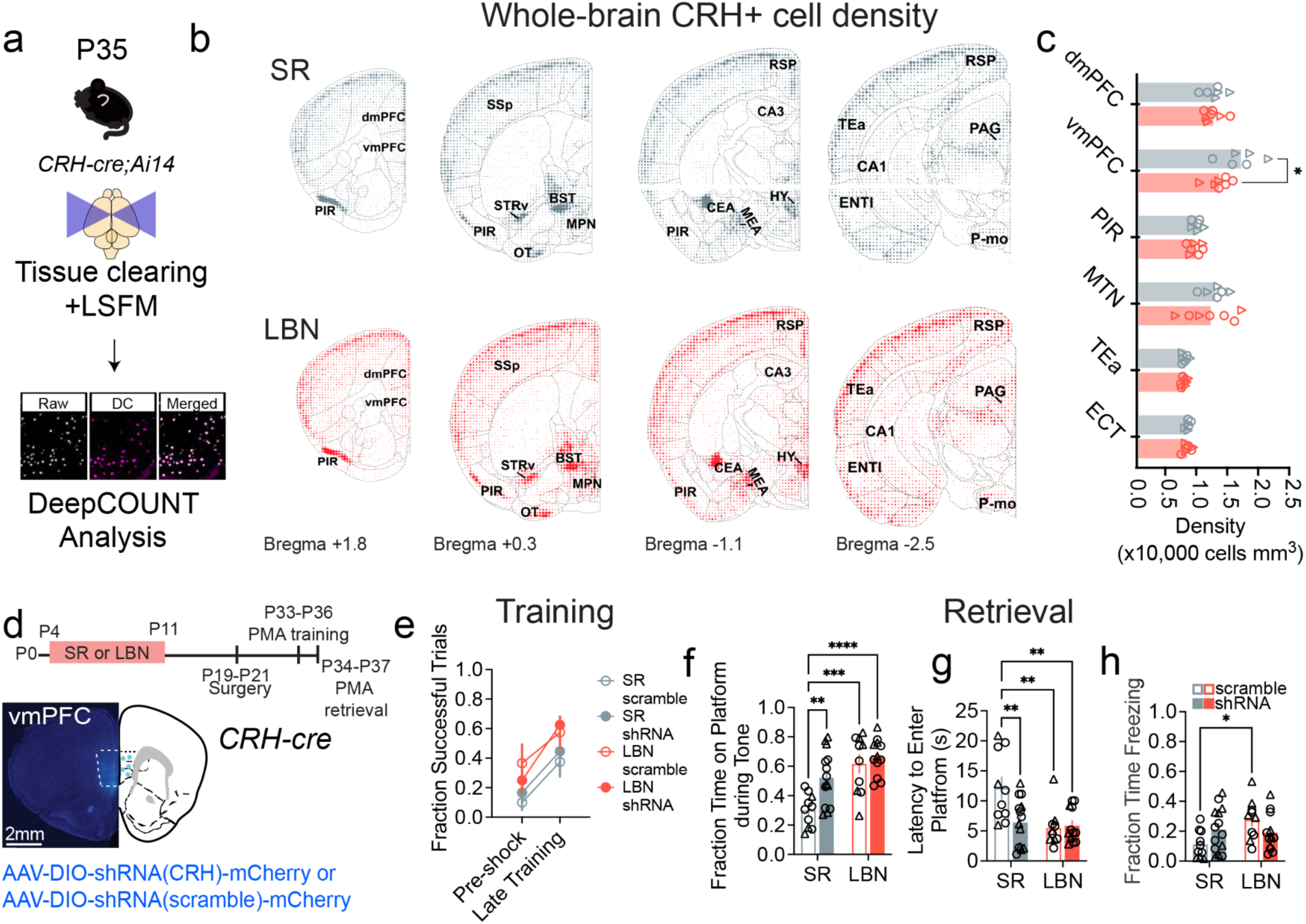
Reduced CRH in the vmPFC is associated with heightened threat avoidance in adolescence. (**a-c**), Analysis of changes in CRH^+^ cell populations in adolescent SR and LBN mice. (**a**) DeepCOUNT analysis pipeline to quantify CRH^+^ cell density across the whole brain (SR, n=6; LBN, n=7 mice). (**b**) Dotograms representing the voxel-wise density of CRH^+^ neurons in SR (gray) and LBN (red) adolescent mice. (**c**) Comparison of CRH^+^ cell density in 6 BLA input regions where retrogradely labelled CRH^+^ cells were present, including dmPFC, vmPFC, PIR, midline group of the dorsal thalamus (MTN), TEa, and ECT (two-way repeated measures ANOVA with Sidak’s post hoc test: region effect *F*_2.18,23.93_ = 41.53, *P*<0.0001; rearing effect *F*_1,11_ = 2.15, *P*=171; interaction *F*_2.18,23.93_ = 2.00, *P* = 0.1544). (**d**) Experimental timeline and image of AAV-shRNA knockdown of Crh expression in the vmPFC (SR: scramble, n=10; shRNA, n=14; LBN: scramble, n=10, shRNA, n=12 mice). (**e**) Fraction of successful trials during PMA training averaged for the pre-shock baseline and the last four tones of the session (three-way ANOVA: trial effect *F*_1,42_=20.58, *P*<0.0001; rearing effect *F*_1,42_=10.16, *P*=0.0027; knockdown effect *F*_1,42_=0.098, *P*=0.756; trial x rearing effect *F*_1,42_=0.013, *P*=0.909; trial x knockdown effect *F*_1,42_=0.47, *P*=0.50; rearing x knockdown effect *F*_1,42_=0.803, *P*=0.375; trial x rearing x knockdown *F*_1,42_=0.417, *P*=0.522). (**f-h**), Behavioral performance during PMA retrieval. (**f**) Fraction of time on the platform averaged across the session (two-way ANOVA with Tukey’s post hoc test: knockdown effect *F*_1,42_=3.37, *P*=0.074; rearing effect *F*_1,42_=22.81, *P*<0.0001; interaction effect *F*_1,42_=7.88, *P*=0.0075). (**g**) Latency to enter the platform following tone onset (two-way ANOVA with Tukey’s post hoc test: knockdown effect *F*_1,42_=9.03, *P*=0.0045; rearing effect *F*_1,42_=4.98, *P*=0.031; interaction effect *F*_1,42_=6.46, *P*=0.015). (**h**) Fraction of time freezing averaged across the session (two-way ANOVA with Tukey’s post hoc test: knockdown effect *F*_1,42_=0.029, *P*=0.867; rearing effect *F*_1,42_=3.60, *P*=0.0646; interaction effect *F*_1,42_=6.06, *P*=0.018). All data points represent biological replicates. Data are shown as mean ± s.e.m., and shading reflects between-subjects s.e.m.; \**P*<0.05,\*\**P*<0.01, \*\*\**P*<0.001, ****P<0.0001 . Triangles denote females and circles denote males.

To determine whether CRH signaling in the vmPFC is involved in threat avoidance behaviors we utilized short-hairpin RNA (shRNA) to knock down *Crh* expression locally (Fig. 2d). The density of CRH+ neurons in the vmPFC increases dramatically between P7 and P21, particularly in layer 2/3, which are the main source of BLA-projecting neurons^56^. We therefore infused our shRNA at ∼P19 to coincide with this developmental surge. We verified knockdown by using *in situ* hybridization to quantify *Crh* mRNA expression. vmPFC^CRH+^ cells co-expressing the anti-*Crh* shRNA and mCherry showed a significant decrease in the intensity of *Crh* labeling compared to mCherry+ cells expressing the scrambled control (Extended Data Fig. 3a). We verified viral regional targeting with histology in each mouse (Extended Data Fig. 3b).

Reducing *Crh* expression in the vmPFC beginning in juvenile stages did not impact the ability of adolescent mice to learn PMA (Fig. 2e). However, vmPFC-specific *Crh* knockdown in SR controls led to a significant increase in time on platform and a decrease in latency to enter the platform during the tone without affecting freezing (Fig. 2f-h). These results indicate that a developmental reduction in vmPFC CRH levels phenocopies ELA-like behavior in adolescence. We observed no changes in avoidance during retrieval in the LBN group (Fig. 2f-h), suggesting that LBN may occlude the effects of *Crh* knockdown by engaging the same mechanism. LBN mice tended to freeze more than controls, but this was not significantly impacted by knockdown of *Crh* (Fig. 1h). Importantly, developmental *Crh* knockdown did not impact anxiety-like behavior in the elevated plus maze or open field assays (Extended Data Fig. 3c-j), highlighting the role of CRH in regulating learned rather than innate threat avoidance. Together with our brain-wide analysis of CRH populations (Fig. 2), these results suggest that an ELA-induced developmental reduction in vmPFC CRH expression contributes to heightened threat avoidance in adolescence.

### ELA enhances activation of a long-range vmPFC^CRH+^→BLA pathway during avoidance

Many CRH expressing cells in the vmPFC are local interneurons that likely do not directly modulate activity in long-range projection targets^30^. Thus, to determine whether there are BLA projecting long-range CRH+ neurons, we injected a retrograde AAV expressing Cre-dependent GFP into the BLA of *CRH-Cre* mice, resulting in fluorescent labeling of CRH^+^ cells that innervate the BLA (Fig. 3a-c, Extended Data Fig. 4a). We observed CRH^+^ BLA-projecting cells in the dorsomedial PFC (dmPFC), piriform cortex (PIR), ectorhinal cortex (ECT) and temporal association areas (TEa), and particularly in the paraventricular nucleus of the thalamus (PVT) and vmPFC (Fig. 3d).

**Figure 3.**
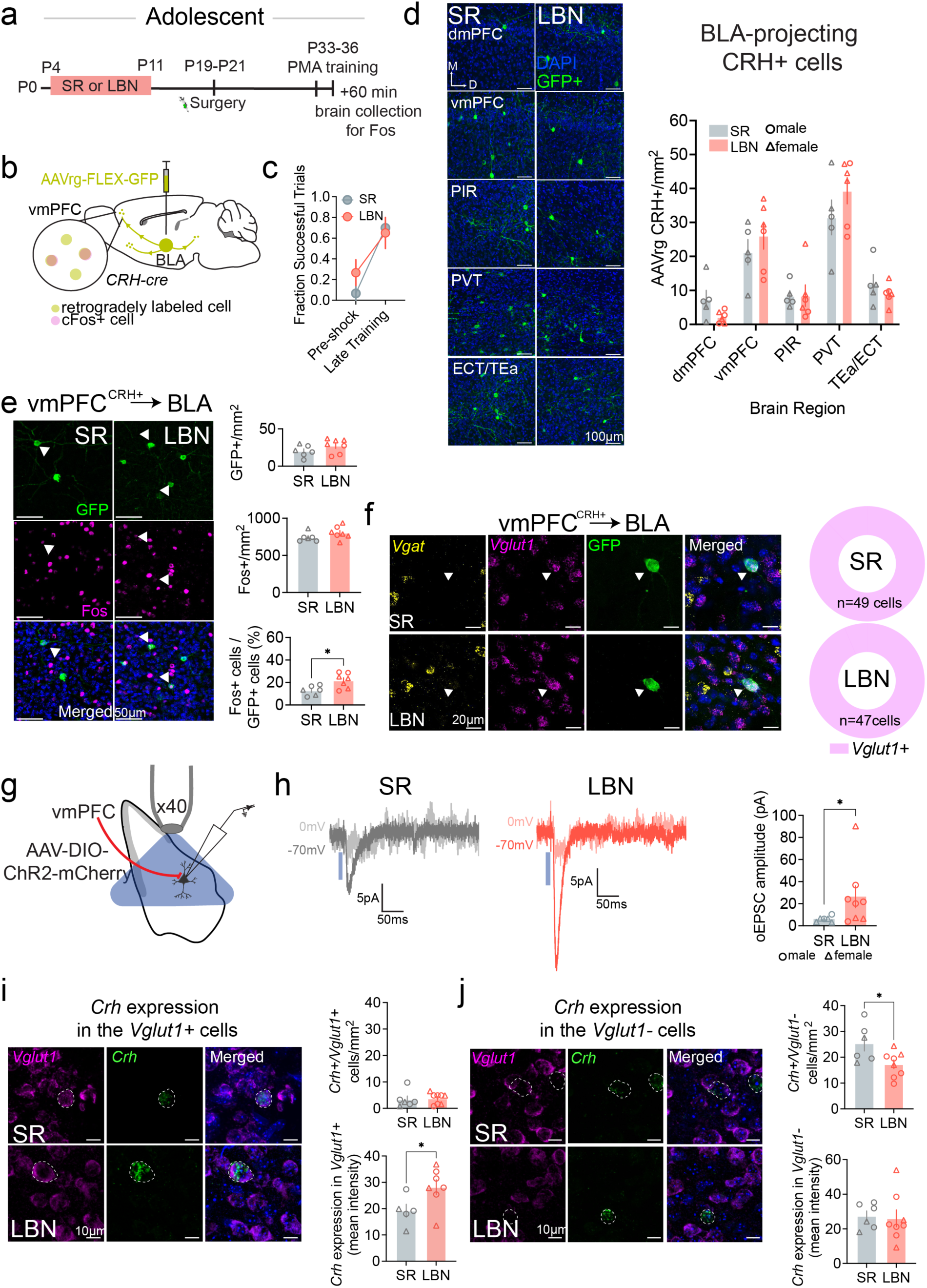
ELA strengthens a vmPFC^CRH+^→BLA pathway. (**a**) Experimental timeline for determining number of activated CRH^+^ cells that project to the BLA during PMA training in adolescent mice. (**b**) Schematic of viral strategy to label CRH^+^ cells that project to the BLA and assess cFos expression. (**c**) Fraction of successful trials during PMA training (SR, n= 6, LBN, n=7 mice) (two-way ANOVA: trial effect *F*_1,11_=30.09, *P*=0.0002; rearing effect *F*_1,12_=0.09, *P*=0.77; interaction effect *F*_1,11_=1.45, *P*=0.254). (**d**) Example images and quantification of retrogradely labelled CRH+ that project to the BLA in the dorsomedial PFC (dmPFC), vmPFC, piriform cortex (PIR), paraventricular nucleus of the thalamus (PVT), ectorhinal cortex (ECT) and temporal association area (TEa) (two-way ANOVA: region effect F2.46,22.2=48.19, p<0.0001; rearing effect F1,9=0.092, p=0.769; interaction effect F2.46,22.2=2.18, p=0.128). (**e**) Retrogradely labelled CRH^+^ cells (green), Fos+ cells (magenta) and merged images in the vmPFC and quantification of the number of GFP^+^ cells (unpaired *t*-test *F*_6,5_=1.47, *P*=0.175), Fos+ cells (unpaired t-test *F*_6,5_=1.91, *P*=0.177), and the percent of GFP^+^/Fos+ cells (unpaired *t*-test *F*_6,5_=2.39, *P*=0.016). (**f**) Examples Vgat+ (yellow), Vglut+ (magenta), vmPFC^CRH+^→BLA cells (green) and merged image in SR and LBN mice and quantification of the percent of vmPFC^CRH+^→BLA cells that are only Vglut+ (magenta) pooled across SR and LBN mice (SR, n=34 cells/5 mice; LBN, n=62 cells/7 mice). (**g,h**), Physiological interrogation of synaptic connectivity of vmPFC^CRH+^→BLA pathway (SR, n=6 cells/4 mice; LBN, n=8 cells/5 mice). (**g**) Schematic of AAV-DIO-ChR2 infection into the CRH^+^ vmPFC cells and patch clamp recording configuration in BLA. (**h**) Representative vmPFC^CRH+^-evoked EPSCs (Vm = 70 mV) and IPSCs (Vm = 0 mV) recorded in BLA principal cells of SR and LBN mice and quantification of optogenetically-evoked excitatory postsynaptic current (oEPSC) amplitude (Welch’s *t*-test *F*_7,5_=5.38, *P*=0.0220). (**i**) Example images of Crh (green) and Vglut1 (magenta) expression in the vmPFC of SR and LBN mice (SR, n= 5, LBN, n=7 mice) and quantification of the number of CRH+ cells that also express Vglut1+ (unpaired *t*-test *F*_5,7_=2.41, *P*=0.800) and the intensity of Crh expression in Vglut1+ cells (unpaired *t*-test *F*_6,4_=1.77, *P*=0.049). (**j**) Example images of Crh (green) and Vglut1 (magenta) expression in the vmPFC of SR and LBN mice and quantification of the number of CRH+ cells that do not express Vglut1 (unpaired *t*-test *F*_5,7_=2.14, *P*=0.029) and the intensity of Crh expression in Vglut1-cells (unpaired *t*-test *F*_5,7_=3.67, *P*=0.836). All data points represent biological replicates. Data are shown as mean ± s.e.m.; \**P*<0.05. Triangles denote females, and circles denote males.

Next, we investigated whether ELA alters the activation of BLA-projecting vmPFC ^CRH+^ neurons during PMA. We measured expression of the immediate early gene Fos, a marker of recently activated neurons, in retrogradely labeled vmPFC^CRH+^→BLA neurons following PMA training (Fig. 3e). The majority of retrogradely labeled vmPFC CRH^+^ cells were located in layer 2/3 (Extended Data Fig. 4b,c), consistent with the layer distribution of the general population of BLA-projecting vmPFC cells^57,58^. We observed no difference in the number of retrogradely labelled GFP+ cells or overall Fos expression within the vmPFC (Fig. 3e). However, in LBN mice, a significantly greater proportion of the GFP+ vmPFC^CRH+^→BLA neurons expressed Fos, indicating they were more activated during PMA training compared to SR controls (Fig. 3e). We observed no stress induced differences in PMA-driven Fos expression in the other BLA-projecting CRH+ populations we identified (Extended Data Fig. 5a-d). Together, these findings suggest that elevated activity in the vmPFC^CRH+^→BLA circuit may contribute to increased threat avoidance in adolescent mice who experience ELA.

### ELA strengthens a glutamatergic vmPFC ^CRH+^→BLA pathway

Reports of long-range CRH+ cell populations are limited, with almost all reporting co-expression with the inhibitory neurotransmitter gamma-aminobutyric acid (GABA)^36,37,56,59^. To determine whether vmPFC^CRH+^→BLA cells are primarily glutamatergic or GABAergic, we combined retrograde viral labeling with *in situ* hybridization to examine expression of the vesicular glutamate transporter 1 (*Vglut1*) and the vesicular GABA transporter (*Vgat*) mRNAs (Fig. 3f). Across groups, all virally labeled neurons expressed *Vglut1*, thus indicating that vmPFC^CRH+^→BLA projection neurons are glutamatergic (Fig. 3f).

We then used channelrhodopsin (ChR2)-assisted circuit mapping^60^ to confirm that the vmPFC^CRH+^→BLA pathway is glutamatergic, and to determine whether LBN affects the strength of synaptic transmission. We injected a cre-dependent AAV-ChR2 into the vmPFC of LBN or SR *CRH-Cre* mice. Allowing two weeks for viral expression, we then prepared acute brain slices from both groups during adolescence (P33-36). Using whole-cell patch clamp recordings in BLA principal cells (Fig. 3g), we measured excitatory and inhibitory postsynaptic currents (EPSCs and IPSCs) evoked by optogenetically stimulating vmPFC^CRH+^ axon terminals within the BLA while voltage clamping neurons at -70 mV or 0 mV, respectively (Fig 3h). No IPSCs were observed in either group, but the amplitude of optogeneticall -evoked EPSCs was significantly greater in LBN mice compared to SR controls (Fig. 3h). Histology confirmed accurate viral targeting (Extended Data Fig. 6a). These findings demonstrate that ELA enhances the excitatory drive from vmPFC^CRH+^ inputs onto BLA principal cells.

To more broadly assess how ELA impacts vmPFC *Crh* expression during adolescence, we also quantified the numbers of CRH+ *Vglut1*+ and *Vglut1*-cells, as well as *Crh* expression levels within those cells. LBN did not affect the number of *Crh*-expressing *Vglut1⁺* neurons, but did lead to increased *Crh* expression relative to SR controls (Fig. 3i). In contrast, *Vglut1*⁻ neurons, which are likely comprise predominantly GABAergic interneurons^61^, showed a significant reduction in the number of *Crh*-expressing cells but no change in *Crh* expression levels (Fig. 3j). Together with the reduction in total vmPFC CRH⁺ cells observed in our whole-brain analysis (Fig. 2), these findings suggest that LBN-driven reduction in CRH+ neurons may be specific to the GABAergic neurons. In contrast, increased *Crh* expression in glutamatergic neurons following ELA may contribute to enhanced downstream activity in the BLA, ultimately promoting heightened avoidance during adolescence.

### vmPFC^CRH+^ →BLA drives increased threat avoidance in LBN adolescents

Previous studies showed that following ELA, elevated activity of central amygdala CRH cells promotes heightened threat reactivity and anhedonia in mice with a history of ELA, suggesting that CRH marks key stress-sensitive pathways that drive enduring changes in emotional behaviors^36^. Given that vmPFC^CRH+^→BLA cells display increased Fos expression during PMA training in LBN mice, enhanced glutamatergic neurotransmission, and increased *Crh* expression, we investigated whether this pathway is required for elevated threat avoidance in adolescent LBN mice, and if so, whether its role extends to adulthood. To do so, we injected the cre-dependent inhibitory opsin PdCO^62^ or a control fluorophore into the vmPFC of *CRH-cre* mice and implanted bilateral fibers over the BLA (Fig. 4a). Two weeks later, we administered blue laser (473 nm, continuous) in adolescents or adults to optogenetically inhibit the vmPFC^CRH+^→BLA pathway during PMA training, specifically during the shock period and 13 seconds following its termination (Fig. 4b). This time period was chosen based on our findings that BLA activity is heightened during this time period (Fig. 1m,n). Correct viral expression and fiber placement were confirmed in all animals (Fig. 4c; Extended Data 6b,c).

**Figure 4.**
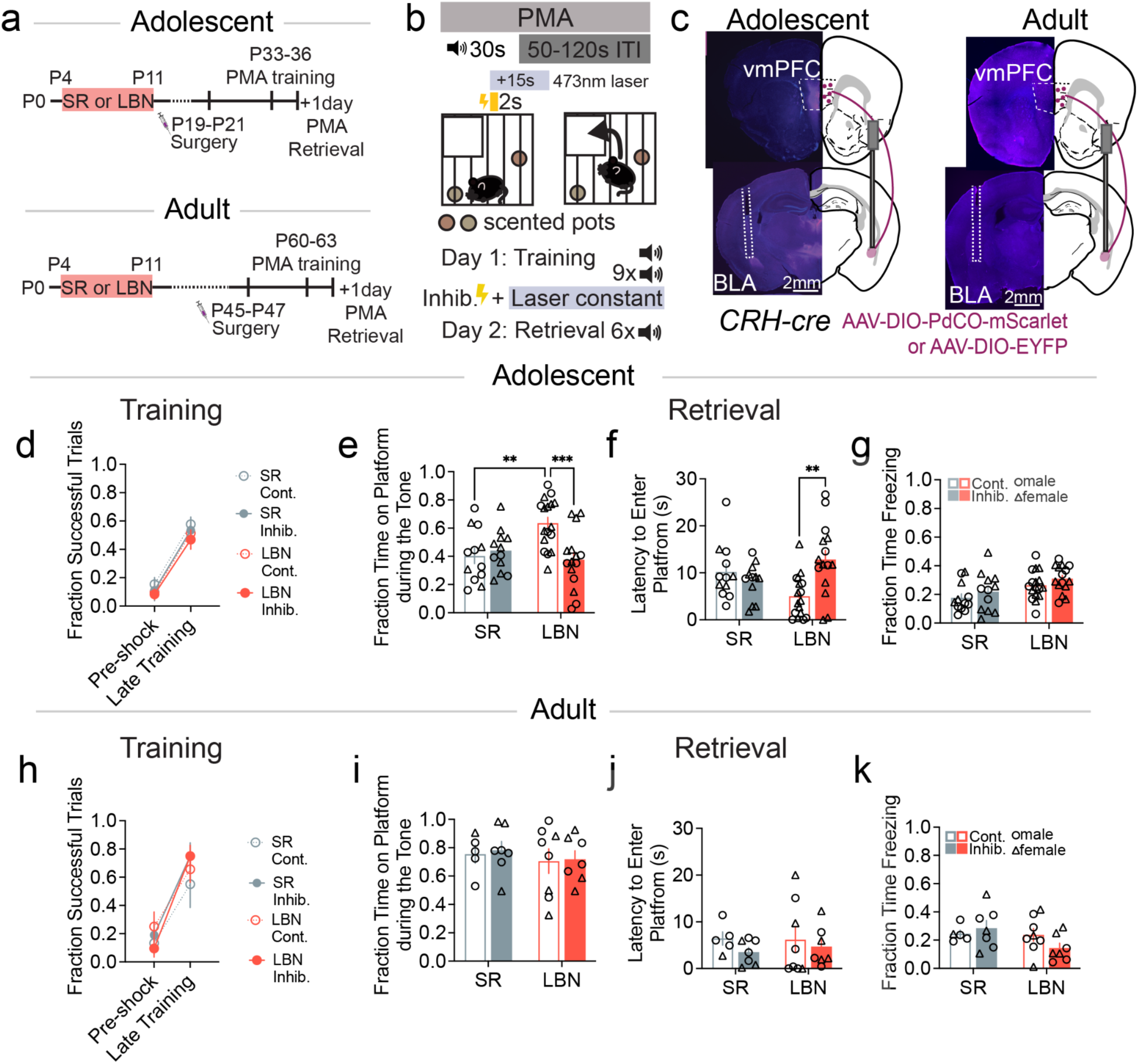
vmPFC^CRH+^→ BLA pathway inhibition reduces ELA-induced avoidance behavior in adolescents, but not in adults. (**a**) Experimental timeline for adolescent and adult cohorts. (**b**) Schematic representation of PMA with constant laser stimulation during the shock period and 13 seconds post shock during training (Adol. SR cont., n=12; SR PdCO n=12; LBN cont., n=15, LBN PdCO, n=17; Adult SR cont., n=5; SR PdCO n=7; LBN cont., n=8, LBN PdCO, n=7 mice). (**c**) Schematic and example images of AAV-DIO-PdCO expression and bilateral fiber optic implants in the BLA of *CRH-Cre* mice. (**d**) Fraction of successful trials in adolescents during PMA training averaged for the pre-shock baseline and the last four tones of the session (three-way ANOVA: trial effect *F*_1,52_=91.51, *P*<0.0001; rearing effect *F*_1,52_=1.74, *P*=0.193; opsin effect *F*_1,52_=0.86, *P*=0.360; trial x rearing effect *F*_1,52_=0.062, *P*=0.804; trial x opsin effect *F*_1,52_=0.028, *P*=0.869; rearing x opsin effect *F*_1,52_=0.0031, *P*=0.956; trial x rearing x opsin *F*_1,52_=0.25, *P*=0.620). (**e-g**), Behavioral performance during PMA retrieval. (**e**) Fraction of time on the platform averaged across the session (two-way ANOVA with Tukey’s post hoc test: rearing effect *F*_1,52_=2.88, *P*=0.10; opsin effect *F*_1,52_=4.98, *P*=0.03; interaction effect *F*_1,52_=8.95, *P*=0.004). (**f**) Latency to enter the platform following tone onset (two-way ANOVA with Tukey’s post hoc test: rearing effect *F*_1,52_=0.029, *P*=0.867; opsin effect *F*_1,52_=3.56, *P*=0.065; interaction effect *F*_1,52_=10.20, p=0.0024). (**g**) Fraction of time freezing during the tone (two-way ANOVA with Tukey’s post hoc test: rearing effect *F*_1,52_=9.47, *P*=0.0033; opsin effect *F*_1,52_=2.33, *P*=0.133; interaction effect *F*_1,52_=0.0026, *P*=0.960). (**h**) Fraction of successful trials in adults during PMA training averaged for the pre-shock baseline and the last four tones of the session (three-way ANOVA: trial effect *F*_1,46_=52.40, *P*<0.0001; rearing effect *F*_1,46_=0.21, *P*=0.652; opsin effect *F*_1,46_=0.49, *P*=0.489; trial x rearing effect *F*_1,46_=0.09, *P*=0.765; trial x opsin effect *F*_1,46_=1.93, *P*=0.171; rearing x opsin effect *F*_1,46_=1.28, *P*=0.264; trial x rearing x opsin *F*_1,46_=0.14, *P*=0.709). (**i-k**), Behavioral performance during PMA retrieval. (**i**) Fraction of time on the platform averaged across the session (two-way ANOVA: rearing effect *F*_1,23_=0.56, *P*=0.460; opsin effect *F*_1,23_=0.07, *P*=0.788; interaction effect *F*_1,23_=0.01, *P*=0.921). (**j**) Latency to enter the platform following tone onset (two-way ANOVA: rearing effect *F*_1,23_=0.052, *P*=0.822; opsin effect *F*_1,23_=1.16, *P*=0.293; interaction effect *F*_1,23_=0.125, *P*=0.27). (**k**) Fraction of time freezing during the tone (two-way ANOVA: rearing effect *F*_1,23_=2.72, *P*=0.113; opsin effect *F*_1,23_=0.356, *P*=0.557; interaction effect *F*_1,23_=2.16, *P*=0.156). All data points represent biological replicates. Data are shown as mean ± s.e.m.; \**P*<0.05, \*\**P*<0.01, \*\*\**P*<0.001. Triangles denote females, and circles denote males.

Inhibition of vmPFC^CRH+^→BLA did not influence the ability of adolescent mice to learn PMA (Fig. 4d). However, during PMA retrieval, in the absence of optogenetic manipulation, those who experienced LBN exhibited a significant decrease in threat avoidance during the shock-predictive tone (Fig. 4e). This difference was driven by an increased latency to enter the safety platform upon tone onset (Fig. 4f) while having no impact on freezing behavior (Fig. 4g). This same manipulation in adults had no impact on either PMA training (Fig. 4h) or retrieval the following day (Fig. 4i-k). Thus, heightened activity of the vmPFC^CRH+^→BLA pathway drives enhanced threat avoidance in the adolescent period, but not later in life. Broad inhibition of the vmPFC→BLA pathway also produced a significant reduction in avoidance behavior in adolescent LBN mice (Extended Data Fig. 7a-g). This indicates that while the broad vmPFC→BLA pathway contributes to heightened threat avoidance in adolescent LBN mice, targeting the CRH^+^ subset alone is sufficient to drive this effect.

We next investigated whether stimulating the vmPFC^CRH+^→BLA pathway in SR controls could induce LBN-like effects on threat avoidance behavior in adolescence. We infused a cre-dependent excitatory opsin into the vmPFC and implanted bilateral fibers over the BLA (Extended Data Fig. 8a,b). We delivered blue laser pulses (473nm, 50ms, 15Hz) to optogenetically stimulate this pathway during PMA training, again during and after the shock period (Extended Data Fig. 8c). We observed no group differences in the ability to learn PMA (Extended Data Fig. 8d) or avoidance levels during retrieval the following day (Extended Data Fig. 8e-g). Together with our finding that this pathway elicits small EPSCs in the BLA in SR mice (Fig. 3k), this suggests that the vmPFC^CRH+^→BLA pathway minimally affects downstream BLA activity and threat avoidance in SR mice. Furthermore, these findings suggest that earlier-occurring developmental circuit plasticity, as opposed to acute increases in pathway activity during adolescence, may contribute to elevated threat avoidance following LBN.

To determine if stimulating or inhibiting the vmPFC^CRH+^→BLA pathway was innately rewarding or aversive, respectively, we performed real-time place preference (RTPP) assay in which one side of a chamber was paired with optogenetic manipulation of this pathway. Activating the vmPFC^CRH+^→BLA pathway produced no significant avoidance of the laser-paired chamber (Extended Data Fig. 8h,i). The same manipulation also had no effect on anxiety-like behavior during a 10-minute open field assay when we interleaved 30 seconds of laser stimulation (Extended Data Fig. 8j-l). Similarly, inhibiting the vmPFC^CRH+^→BLA pathway also had no effect during the RTPP or open field assays in adolescents (Extended Data Fig. 9a-e). We observed largely similar results in adults (Extended Data Fig. 9f-i) except that locomotion was reduced following inhibition in the open field assay (Extended Data Fig. 9j). Taken together these findings indicate that the vmPFC^CRH+^→BLA pathway plays a distinct role in learned threat avoidance behavior and does not drive innate aversion.

### *Crh* expression in vmPFC^VGLUT^^1^ neurons is required for LBN-induced heightened avoidance

Our optogenetic experiments reveal that activity in the vmPFC^CRH^→BLA pathway is required for heightened avoidance in LBN adolescent mice. However, this manipulation may influence both glutamate and CRH release. To determine whether and when CRH signaling is required for LBN-induced increases in adolescent avoidance and associated changes in BLA activity we infused a Cre-dependent shRNA targeting CRH into the vmPFC of *VGLUT1-Cre* mice. This strategy selectively reduces CRH in excitatory vmPFC neurons, which comprise the vmPFC^CRH^→BLA pathway. Adolescent mice received surgery at P19–P21 and adults at P45-47. Two weeks later mice underwent PMA training and retrieval, followed by perfusion to assess Fos expression in the BLA (Fig. 5a). Correct viral expression was confirmed in all animals (Extended Data Fig. 10a,b).

**Figure 5.**
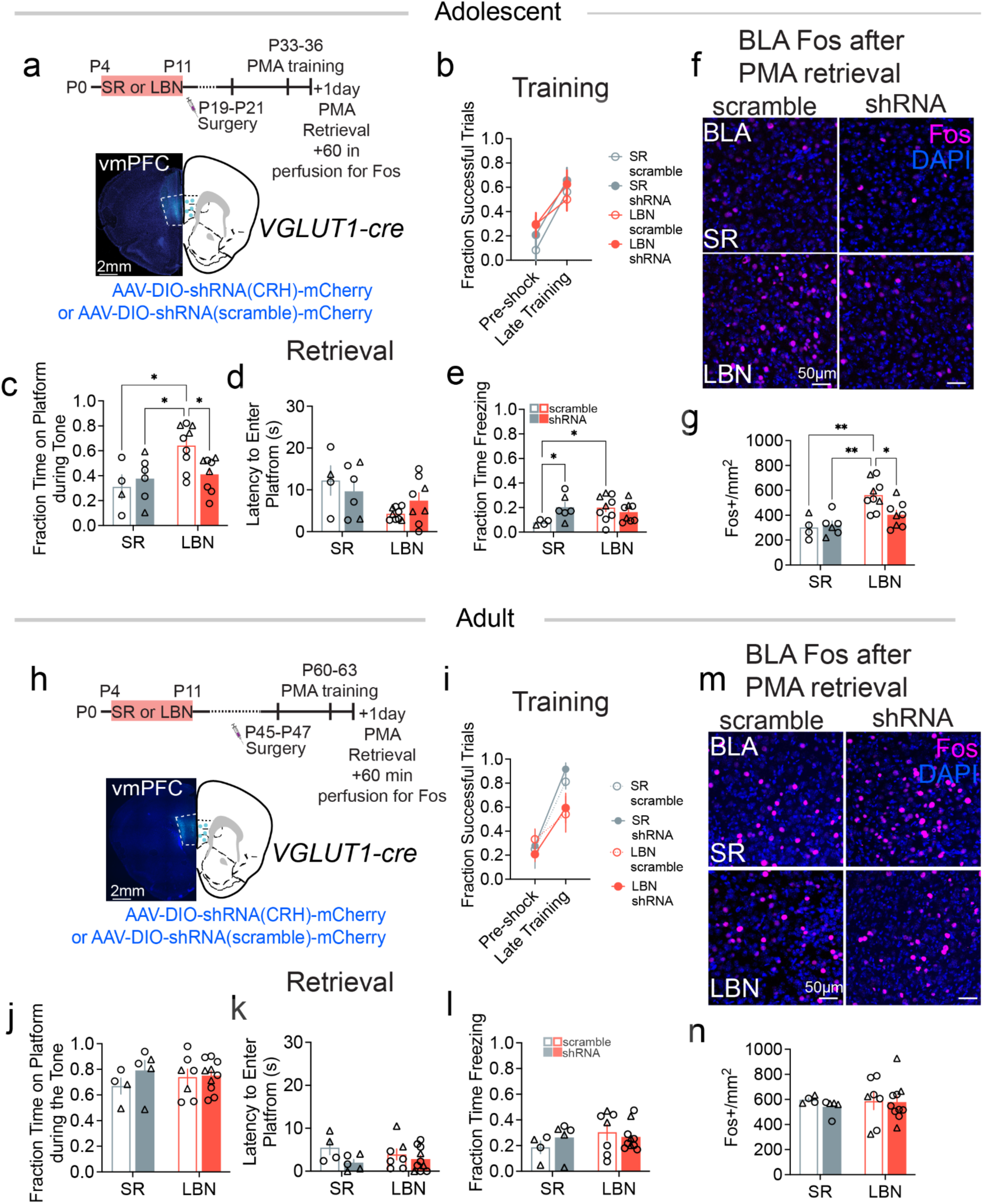
Reduction of *Crh* expression in the vmPFC→ BLA pathway attenuates ELA-induced heightened avoidance and BLA activity in adolescents, but not in adults. (**a**) Experimental timeline and image of AAV-shRNA knockdown of *Crh* expression in vmPFC^VGLUT1+^ cells in adolescents (SR: scramble, n=4; shRNA, n=6; LBN: scramble, n=8, shRNA, n=7 mice). (**b**) Fraction of successful trials during PMA training averaged for the pre-shock baseline and the last four tones of the session (three-way ANOVA: trial effect *F*_1,23_=21.25, *P*=0.0001; rearing effect *F*_1,23_=0.093, *P*=0.764; knockdown effect *F*_1,23_=2.99, *P*=0.097; trial x rearing effect *F*_1,23_=1.41, *P*=0.247; trial x knockdown effect *F*_1,23_=0.067, *P*=0.800; rearing x knockdown effect *F*_1,23_=0.68, *P*=0.417; trial x rearing x knockdown F_1,23_=0.32, p=0.576). (**c-e**), Behavioral performance during PMA retrieval. (**c**) Fraction of time on the platform averaged across the session (two-way ANOVA with Tukey’s post hoc test: knockdown effect *F*_1,23_=1.42, *P*=0.246; rearing effect *F*_1,23_=7.16, *P*=0.014; interaction effect *F*_1,23_=4.68, *P*=0.041). (**d**) Latency to enter the platform following tone onset (two-way ANOVA with Tukey’s post hoc test: knockdown effect *F*_1,23_=0.02, *P*=0.889; rearing effect *F*_1,23_=6.53, *P*=0.018; interaction effect *F*_1,23_=2.06, *P*=0.164). (**e**) Fraction of time freezing averaged across the session (two-way ANOVA: knockdown effect *F*_1,23_=1.48, *P*=0.237; rearing effect *F*_1,23_=1.31, *P*=0.264; interaction effect *F*_1,23_=4.95, *P*=0.036). (**f**) Example images of Fos expression in the BLA following Crh knockdown in vmPFC^VGLUT1+^ cells. (**g**) Quantification of Fos+ cells per mm^2^ (two-way ANOVA with Tukey’s post hoc test: knockdown effect *F*_1,23_=2.69, *P*=0.115; rearing effect *F*_1,23_=16.33, *P*=0.0005; interaction effect *F*_1,23_=4.31 *P*=0.049. (**h**) Experimental timeline and image of AAV-shRNA knockdown of *Crh* expression in vmPFC^VGLUT1+^ cells in adults (SR: scramble, n=4; shRNA, n=5; LBN: scramble, n=7, shRNA, n=10 mice). (**i**) Fraction of successful trials during PMA training averaged for the pre-shock baseline and the last four tones of the session (three-way ANOVA: trial effect *F*_1,40_=30.87, *P*<0.0001; rearing effect *F*_1,40_=3.22, *P*=0.0803; knockdown effect *F*_1,40_=0.033, *P*=0.856; trial x rearing effect *F*_1,40_=3.50, *P*=0.067; trial x knockdown effect *F*_1,40_=0.615, *P*=0.437; rearing x knockdown effect *F*_1,40_=0.40, *P*=0.530; trial x rearing x knockdown *F*_1,40_=0.097, *P*=0.757). (**j-l**), Behavioral performance during PMA retrieval. (**j**) Fraction of time on the platform averaged across the session (two-way ANOVA: knockdown effect *F*_1,22_=1.61, *P*=0.293; rearing effect *F*_1,22_=0.059, *P*=0.810; interaction effect *F*_1,22_=0.82, *P*=0.376). (**k**) Latency to enter the platform following tone onset (two-way ANOVA: knockdown effect *F*_1,22_=3.55, *P*=0.073; rearing effect *F*_1,22_=0.092, *P*=0.765; interaction effect *F*_1,22_=0.099, *P*=0.330). (**l**) Fraction of time freezing averaged across the session (two-way ANOVA: knockdown effect *F*_1,22_=0.15, *P*=0.707; rearing effect *F*_1,22_=1.31, *P*=0.264; interaction effect *F*_1,22_=1.13, *P*=0.300). (**m**) Example images of Fos expression in the BLA following *Crh* knockdown in vmPFC^VGLUT1+^ cells. (**n**) Quantification of Fos+ cells per mm^2^ (two-way ANOVA: knockdown effect *F*_1,22_=0.312, *P*=0.582; rearing effect *F*_1,22_=0.056, *P*=0.815; interaction effect *F*_1,22_=0.186 *P*=0.670. All data points represent biological replicates. Data are shown as mean ± s.e.m.; \**P*<0.05, \*\**P*<0.01, \*\*\**P*<0.001. Triangles denote females, and circles denote males.

Developmental *Crh* knockdown did not impair acquisition of learned avoidance in either SR or LBN adolescent mice (Fig. 5b). During retrieval, *Crh* knockdown had no effect in SR controls but significantly reduced avoidance in LBN mice, as measured by decreased time spent on the platform (Fig. 5c), without altering latency to enter or freezing (Fig. 5d,e). Consistent with these behavioral effects, *Crh* knockdown reduced Fos expression in the BLA (Fig. 5f,g). To determine whether ELA-induced changes in *Crh* expression are important specifically during development, an adult comparison group underwent the same procedures (Fig. 5h). We observed no behavioral differences between SR and LBN adult mice and *Crh* knockdown did not influence PMA behavior (Fig. 5i-l) or BLA Fos expression in either rearing group (Fig. 5m,n). To confirm these changes were specific to learned avoidance we also measured anxiety-like behavior and locomotion using the open field and elevated zero maze. These were assessed prior to PMA training and we observed no effect of *Crh* knockdown in either rearing group (Extended Data Fig. 10c-h). Together, these data indicate that juvenile and adolescent stages represent a key developmental window when alterations in vmPFC CRH signaling can influence threat avoidance circuit function.

## DISCUSSION

ELA is a major risk factor for developing mental illness, including anxiety and depression, which are characterized by high levels of avoidance and often manifest during adolescence^2,3,7–12^. Despite this, little is known about how ELA shapes the maturation of threat avoidance circuits during this critical developmental window. ELA can take many forms, and both the nature and the timing of adversity impact behavioral outcomes^63^. In rodent models, it is widely accepted that chronic stress within the first ten days of life has enduring consequences, though different models offer unique advantages^45^. Here we selected the LBN model because it requires minimal experimenter intervention and captures common human experiences in which the mother is present but provides fragmented and unpredictable care^45^. Using this model, we identified a long-range, glutamatergic vmPFC^CRH+^→BLA pathway that, following ELA, displays enhanced activation during threat avoidance, synaptic strengthening, and increased *Crh* expression during adolescence. Inhibiting the vmPFC^CRH+^→BLA pathway during behavior or reducing vmPFC *Crh* expression in exclusively excitatory cells earlier in development rescues ELA-induced elevations in BLA activity and threat avoidance in adolescence, but the same manipulations did not affect threat avoidance in adulthood. Interestingly, ELA also reduces the overall density of CRH⁺ cells in the vmPFC. In line with this, we find that developmental, vmPFC-wide *Crh* knockdown in SR mice phenocopies the heightened avoidance induced by ELA. Together, these findings suggest that in animals that experienced ELA, CRH acts in a cell type-specific manner to induce developmental plasticity in vmPFC circuits critical for threat responding. By revealing a previously unrecognized, molecularly defined prefrontal circuit that is sensitive to ELA and capable of driving heightened threat avoidance during adolescence, our findings reveal a potential circuit mechanism through which ELA may contribute to vulnerability to psychiatric illness.

Previous work has shown that individuals exposed to ELA exhibit heightened BLA activation following exposure to fearful stimuli, as measured by functional magnetic resonance imaging in humans and Fos labeling in rodents ^20,64,65^. Consistent with this, we find that adolescent mice exposed to ELA have increased activity in the BLA (Fig. 1). During PMA, BLA activity is elevated during the shock experience and the conditioned tone in LBN animals compared to controls. PMA uses a lower shock intensity compared to classical fear conditioning, which may explain why SR controls have relatively low levels of tone-evoked activity^66^. Together, these findings suggest that ELA sensitizes the BLA to even weakly aversive stimuli during learning, driving elevated responses to threat-predictive cues.

CRH is a stress-responsive neuropeptide that promotes fear learning and enhances neuronal excitability, suggesting it may mediate ELA-induced changes in mPFC-BLA circuits^49–51^. Although altered CRH expression is evident in the hypothalamus, amygdala and hippocampus, prior work focused on adults^33,34,67–69^. We observed a developmental, vmPFC-specific decrease in *Vglut-*CRH^+^ cells, with no change in the number of *Vglut+* CRH cells, including vmPFC^CRH+^→BLA projectors (Fig. 3). Most vmPFC CRH^+^ neurons are GABAergic interneurons that co-express the vasoactive intestinal peptide (VIP)^30^, though studies have noted sparse long-range GABAergic CRH^+^ projections to regions such as the nucleus accumbens and lateral septum^36,37,56^. Thus, the reduction in CRH^+^ cells observed in our whole-brain analysis and *in situ* data likely reflects a loss of *Crh* expression in vmPFC GABAergic neurons. Furthermore, our vmPFC-wide CRH knockdown experiment, which phenocopied the effects of ELA in SR mice (Fig. 2), likely impacted those cells most heavily. VIP interneurons preferentially target somatostatin (SST) interneurons, thereby relieving SST-mediated dendritic inhibition of pyramidal neurons and increasing excitatory output^70,71^. In addition to this synaptic interaction, a subset of SST interneurons expresses CRH receptor 2. These findings raise the possibility that CRH release from VIP interneurons may modulate prefrontal microcircuit maturation^72,73^. While the functional consequences of this reduction remain unclear, such changes could ultimately influence prefrontal output to downstream regions, including the basolateral amygdala.

We also found that ELA-driven plasticity in the vmPFC^CRH+^→BLA circuit promoted heightened threat avoidance during adolescence but had no effect in adulthood. During typical development, reduced threat response during this period may facilitate risky exploration that is important for navigating the environment and establishing adaptive behavioral strategies^11,74^. ELA-induced enhancement of vmPFC^CRH+^→BLA activity may therefore impose a persistent avoidance bias, potentially restricting behavioral repertoires and limiting acquisition of flexible strategies for responding to the environment. Our previous research has shown that, in mice raised under typical conditions, higher levels of threat avoidance observed in adulthood is due to heightened activity, synaptic strength and connectivity within the dmPFC to BLA pathway^40^. Here we similarly observed high avoidance in adults, regardless of rearing condition. Further, inhibition of vmPFC^CRH+^→BLA or knockdown of *Crh* expression in excitatory vmPFC neurons did not significantly alter avoidance at this age, suggesting that this pathway has a more potent role during adolescence. It is possible that other circuits take on a larger role in modulating PMA behavior by adulthood, diminishing or masking the contribution of the vmPFC^CRH+^→BLA. Overall, the selective effects of vmPFC-BLA inhibition and pathway-specific CRH knockdown during adolescence highlights this circuit as a potential target for early intervention. Together, by linking mPFC circuit development to the enduring effects of ELA, we can provide a mechanistic framework for understanding how early-life experiences shape neurodevelopmental trajectories and ultimately influence vulnerability to psychiatric disorders.

## ACKNOWLEDGMENTS

We thank the laboratories Drs. Laura DeNardo and Scott Wilke for project discussion. We thank Chloe Christensen for assistance with mouse colony management. We thank Russell Ahmed for assistance with perfusions. We thank Dr. Carlos Portera-Cailliau, Dr. Weizhe Hong, and Abigail Yu for helpful comments on the manuscript. We acknowledge the Broad Stem Cell Research Center Microscopy Core at the University of California, Los Angeles, for providing access to microscopes.

## FUNDING

This work was funded by the National Institutes of Health grant F31MH138135 (CMG), National Science Foundation Graduate Research Fellowship (CMG), National Institutes of Health grant F30MH134633 (MWG), National Institutes of Health grant R01MH127214 (LAD) and Vallee Foundation Scholars Award (LAD).

## AUTHOR CONTRIBUTIONS

CMG: conceptualization, data collection, data analysis, visualization, manuscript writing and editing. MWG: data collection, data analysis, visualization, manuscript editing. ZZ, CB, MS, CBK, NSJ, MR, MA, RH, JU: data collection and analysis. LAD: conceptualization, data collection, data analysis, visualization, manuscript writing and editing, supervision, funding acquisition, projection administration.

## COMPETING INTERESTS

Authors declare that they have no competing interests.

## MATERIALS AND CORRESPONDENCE

Requests for further information and resources should be directed to and will be fulfilled by the lead contact, Laura DeNardo

## METHODS

### Subjects

Female and male C57B16/J (JAX Stock No. 000664), *CRH-Cre* (JAX Stock No. 012704), *VGLUT1-Cre* (JAX Stock No. 023527) and Ai14 tdTomato Cre-reporter (JAX Stock No. 007914) transgenic mice were bred in house. Post weaning mice were group-housed (2-4 per cage). Experimental endpoints occurred during mid-adolescence, postnatal day (P)33-37. Mice were kept on a 12 hr light cycle (lights on 7am-7pm) in a temperature and humidity-controlled room. Food and water were available ad libitum. All procedures followed animal care guidelines approved by the University of California, Los Angeles Chancellor’s Animal Research Committee.

### Limited Bedding and Nesting Protocol

Dams underwent ELA using a limited bedding and nesting protocol (LBN), a model of resource scarcity in which dam and pups were placed in low bedding conditions with limited access to nesting material for 7 consecutive days (P4-P11). This model has been previously validated to disrupt maternal care^45^. On P4, dam and pups were transferred from their standard home cage, which included two 1-inch x 1-inch cotton nestlet and a wood-pulp disposable hut, to a LBN cage containing a wire mesh floor and one 1-inch x 1-inch cotton nestlet. Standard reared control dam and pups were moved into a new standard home cage. All mice had continued access to food and water. Following one week (P11), pups and dams were returned to their standard housing with full bedding and nesting material. Maternal behavior was recorded for 1 hour from 8am-9am and 4pm-5pm every day for a week. Maternal behavior was assessed by manually measuring the proportion of time the dam was on the nest during the morning recording of P6. All pups were weaned at P21.

### Surgery

At P19-21 mice were induced in 3% isoflurane in oxygen until loss of righting reflex and transferred to a stereotaxic apparatus. The stereotax was fitted with an attachment for developing mice including a small bite bar and developmental ear bars. A nonsteroidal anti-inflammatory agent was administered pre- and postoperatively to minimize pain and discomfort. The mouse’s head was shaved and prepped with three scrubs of alternating betadine and then 70% ethanol. 2% Lidocaine was injected under the scalp as a local anesthetic. A small incision was made in the scalp. For viral injections, a small hole was drilled above the injection target and a hamilton syringe loaded with virus was lowered to the correct stereotaxic coordinate. Virus was pressure injected at 100nL/minute, and following completion of the injection the syringe was left in place for 5 minutes and then slowly removed from the skull. For all experiments animals were allowed to recover and then placed back into a clean homecage with their mother and littermates until P21, when they were weaned. Animals were excluded if virus expression or fiber placement was mistargeted.

#### Fiber photometry, optogenetics and CRH knockdown

For photometry experiments mice were infused unilaterally with an AAV expressing the genetically encoded calcium indicator GCaMP6s (AAV9-CAG-GCaMP6s-WPRE-SV40, Addgene# 100844, 300 nl) in the BLA (anterior-posterior(AP): -1.6, medial-dorsal(ML): -3.20, dorsal-ventral(DV): -4.8). For optogenetic experiments mice were bilaterally infused with AAVs (inhibition: AAV5-Ef1a-DIO-PdCO-mScarlet, Addgene# 198516, or excitation: AAV8-nEF-Con/Foff 1.0-ChR2-EYFP, Addgene# 137163, 200nl) or control AAVs (inhibition: AAV5-Ef1a-DIO-EYFP, Addgene 27056, excitation: AAV8-Ef1a-DIO-mCherry, Addgene# 114471, 200nl) into the vmPFC (AP 1.8, ML ±0.25, DV -2.9). For broad vmPFC→BLA inhibition AAV5-hysn-PdCO-EGFP (Addgene# 198513, 200nl) and AAV5-hysn-mCherry (Addgene# 114472, 200nl) were used. To accommodate for skull growth, unilateral (photometry, 400 µm, 0.50 NA; Thor Labs) or bilateral (optogenetics, 200µm, Newdoon) optic fibers were implanted in a separate surgery 5 days prior to PMA training and secured with Metabond (Parkell). For knockdown experiments mice were infused bilaterally with a cre-dependent AAV expressing anti-CRH shRNA (AAV2/9-CMV-DIO-(mCherry-U6)-shRNA(CRH)-WPRE-hGHpA, BrainVTA, PT-12461, 200nl) or a scrambled control (AAV2/9-CMV-DIO-(mCherry-U6)-shRNA(scramble)-WPRE-hGHpA, BrainVTA, PT–2788, 200nl).

#### Viral tracing

Mice were infused unilaterally infused with 150nl mixture of a 3:1 dilution of a cre-dependent AAV (AAVrg-CAG-FLEX-GFP, Addgene# 51502) and non-cre dependent AAV (AAV5-CAG-tdTomato, Addgene# 58462) in the BLA (AP -1.6, ML -3.20, DV -4.8). The latter was used to verify successful targeting of the retrograde virus; animals were excluded if expression was not in the BLA.

### Behavioral Assays

#### Platform-Mediated Avoidance

Mice were handled for 3 days preceding the behavioral testing procedure. The conditioning chamber consisted of an 18 x 30 cm cage with a grid floor wired to a scrambled shock generator (Lafayette Instruments). The chamber was surrounded by a custom-built acoustic chamber and scented with 50% Windex. A thin acrylic platform (1.3 cm thick) covered 25% of the floor. Two small weigh boats filled with vanilla or almond extract were placed beneath the floor to encourage exploration of the chamber by the mice. At mid-adolescence (P33-36) training occurred and mice were presented with three baseline 30s 4 kHz tones (CS), followed by nine presentations of the CS that co-terminated with a 2 s foot shock (0.13mA). Mice were perfused either 10 minutes (pCREB expression) or 1 hour (Fos expression) after training and their brains collected and processed for Fos immunostaining.

For all other experiments, mice were presented with six CS in the absence of shocks the following day to probe ability to retrieve and express avoidance memory. Tones were separated by randomized interval lengths that ranged from 80 to 150 seconds. For CRH knockdown in vmPFC^VGLUT1+^ cells mice were perfused 1 hour following training for Fos expression.

#### Shock Sensitivity

To assess the minimum foot-shock intensity required to elicit a behavioral response (vocalization, scurry or dart), naive adolescent mice were placed in the same operant conditioning chamber as in PMA, but without the platform. Mice were exposed to a series of foot shocks, beginning at 0.02 mA and increased at 0.02 intervals until 0.20 mA. The amplitude of the foot-shock at which a given mouse first vocalized, scurried and darted was recorded. Vocalization was defined as the emittance of an audible sound. Scurry was defined as rapid stepping with the absence of jumping. Dart was defined as a high velocity, horizontal jump.

#### Open-field Test

Mice were acclimated to the testing room for 10 minutes and then placed in a plastic arena (50 x 50 x 40 cm). Locomotor activity and time spent in the center of the arena were recorded during a 5 (naive mice) or 10 (optogenetic and knockdown manipulations) minute test using a webcam. The video-tracking system, BioViewer, was used to analyze the data (naïve) while EZTrack^75^ was used to analyze the remaining datasets. The arena was divided into two zones, the center (25% of the total area) and periphery (75% of the total area) and time spent and entries in each zone, as well as total distance traveled, was recorded. For optogenetic experiments, recordings began with a 60-s baseline period, followed by four cycles of 30-s laser on and 30-s laser off. During these periods, time spent in each zone, number of zone entries, and total distance traveled were quantified and compared between laser on and off epochs.

#### Elevated Plus Maze

Mice were acclimated to the testing room for 10 minutes. Mice were placed into the center of the plus-shaped maze with their nose pointing towards an open arm. Locomotor activity and time spent in the open and closed arms and center of the arena were recorded for 5 minutes using a webcam. Bioviewer was used to analyze the data by dividing the arena into two sets of open arms, two of closed arms and the center. The time spent and entries in each zone, as well as total distance traveled, was recorded.

#### Elevated Zero Maze

Mice were acclimated to the testing room for 10 minutes. Mice were placed in a closed arm with the nose pointing towards an open arm of a zero-shaped elevated maze. Locomotor activity and time spent in the open and closed arms and center of the arena were recorded for 10 minutes using a webcam. EZTrack was used to analyze the data. The time spent and entries in each zone, as well as total distance traveled, was recorded.

#### Novel Odor Assay

On day 1, mice were placed in an empty plastic arena (50 x 50 x 40 cm) for 10 minutes of habituation. On day 2, mice were placed back in the arena with two identical scented plastic spheres (same scent for each: almond or vanilla, counterbalanced across mice). Mice were left in the chamber until they interacted with the spheres for a total of 20s. Mice were excluded if they did not reach this criterion within 10 minutes. Interaction was defined as being within 2 cm from the sphere with the nose pointed at it. On day 3, mice were placed in the arena for 10 min and one sphere was scented with the original scent and the other was scented with a novel odor (sides counterbalanced between animals). Interaction time was quantified from video recordings. Groups represent pooled results from multiple, independently run behavioral cohorts.

### Brain-clearing and whole-brain imaging

Mouse brain tissue was prepared following a modified version of the Adipo-Clear Protocol^76^. Briefly, mice were intracardially perfused on ice with 20mL phosphate-buffered saline (PBS, Invitrogen) followed by 4% paraformaldehyde (PFA; Electron Microscopy Sciences). Brains were then hemisected approximately 1mm lateral to the midline and post-fixed overnight at 4℃ in 4% PFA. The following day, samples were sequentially dehydrated using methanol (MeOH, Fisher Scientific) mixed with B1n buffer (1:1000 Triton X-100, 2% w/v glycine, 1:10,000 NaOH 10N, 0.02% sodium azide)/Each methanol gradient (20%, 40%, 60%, and 80%) was applied for 1 hour using a nutator (VWR). Samples were then rinsed twice with 100% MeOH for 1 hour each and subsequently incubated overnight in a 2:1 dichloromethane (DCM) solution. The next day, samples were washed twice in 100% DCM for 1 hour each, followed by three washes in 100% MeOH for progressively longer durations (30 minutes, 45 minutes, 1 hour). Tissue samples were then bleached for 4 hours in a 5:1 H_2_O_2_ solution. Rehydration was achieved through a series of MeOH/B1n buffer washes in decreasing methanol concentrations (80%, 60%, 40%, and 20%) for 30 minutes each, followed by a final 1-hour wash in B1n buffer. Permeabilization was carried out with 5% DMSO/0,3 M glycine in PTxWH buffer for 1 hour, followed by an additional 2-hour incubation in fresh permeabilization solution. Samples were then rinsed in PtxWH for 30 minutes and left in fresh PtxWH buffer overnight. On the following day, two more PtxWH washes (1 hour and 3 hours) were performed.

Samples were incubated with a primary anti-RFP antibody (Rockland Immunochemicals) at a dilution of 1:300 in PTxWH, with continuous shaking at 37℃ for 11 days. This was followed by sequential PTxWH washes: twice for 1 hour each and twice for 2 hours each, with additional PTxWH exchanges over a 2-day period at 37℃. Following primary antibody incubation, samples were incubated with secondary antibody at 1:300 (AlexaFluor 647, ThermoFisher Scientific) at 37℃ for 8 days, with regular PTxWH washes over two days. After antibody staining, samples were rinsed in PBS twice (1 hour each), then for 2 hours twice, and left overnight, Dehydration involved a graded series of MeOH washes (205, 40%, 60%, and 80%) for 30 minutes each, followed by three 100% MeOH washes (30 minutes, 1 hour, and 1.5 hours). Samples were incubated overnight in a 2:1 DCM solution on a nutator. The next day, samples were washed twice in 100% DCM for 1 hour each, then cleared in 100% dibenzyl ether (DBE), with DBE refreshed after 4 hours. Cleared samples were stored in DBE at room temperature in darkness, and imaging was performed after at least 24 hours.

#### Whole-Brain Imaging

Brain samples were imaged using a light-sheet microscope (Ultramicroscope II, LaVision Biotec) outfitted with a sCMOS camera (Andor Neo) and a 2x/0,5 NA objective lens (MVAPLAPO 2x) with a 6mm working distance dipping cap. Image stacks were captured at a 0.8x optical zoom and controlled through Imspector Microscope v285 software. For cell imaging, 488nm and 640nm lasers (20% laser power) were used. Scanning was performed with a 3 µm step size, employing a continuous light-sheet scanning method with a blend algorithm for the 640nm channel (20 acquisitions per plane), and without horizontal scanner for the 488-nm channel.

### Ex vivo electrophysiology

*Surgery* To optogenetically stimulate vmPFC^CRH+^ axons in the BLA, we injected 300nl of AAV8-Ef1a-double floxed-hChR2(H134R)-mCherry)WPRE-HGHpA (Addgene, 20297) into the left vmPFC using techniques described above. Surgeries were performed 16-18 days before recordings.

#### Acute brain slice preparation

To prepare acute brain slices, mice we anesthetized with isoflurane and transcardinally perfused with ice-cold slicing artificial cerebrospinal fluid (ACSF) solution containing 2.5 nM KCL, 1 mM NaH_2_PO_4_, 26.2 mM NaHCO_3_, 4 mM MgCl_2_, 11 mM glucose, 210.3 mM sucrose, 0.5 mM CaCl_2_ and 0.5 mM sodium ascorbate (bubbled with 95% O_2_/5% CO_2_). The brain was rapidly dissected, and 300 µm BLA sections were obtained from the hemisphere ipsilateral to the injection site using a Leica VT1200S vibratome. Slices were transferred to normal ACSF containing 125 nM NaCL, 2,5 nM KCl, 26.2 mM NaHCO_3_, 2 mM MgCl_2_, 11 mM glucose and 2 mM CaCl_2_ (bubbled with 95% O_2_/5% CO_2_) and held at 34 ℃ for 34-40 minutes. Slices were then allowed to cool to room temperature. Slices containing the vmPFC were also collected to verify the injection site.

#### Slice electrophysiology

Recordings were performed at room temperature in normal ACSF. The BLA was identified using white matter tracts and vmPFC^CRH+^ axon fluorescence. Cells were visualized under infrared-differential interference contrast through a x40 objective. Voltage clamp experiments were performed using borosilicate pipettes (5-7 MΩ) filled with internal solution containing 117 mM cesium methanesulfonate, 20 mM HEPES, 0,4 mM EGTA, 2.8 mM NaCl, 5 mM TEA-Cl, 4 mM Na_2_-ATP and 0.4 Na-GTP, adjusted to pH 7.3 using cesium hydroxide (280-290 mOsm). Spontaneous excitatory and inhibitory currents were recorded via 3 minute gap free recordings while holding neurons at -70mV or 0mV, respectively. Excitatory currents from vmPFC^CRH+^ terminal stimulation were obtained by holding neurons at –70 mV and delivering 0.5 ms of 50-mW blue (∼470-nm) light through a ×40 objective using a CoolLED pE-300Ultra light source. Paired-pulse recordings were performed by delivering identical light stimuli spaced 100ms apart. Only neurons with consistent synaptic responses to optical stimulation were included in the evoked current dataset. Inhibitory currents were recorded in the same way but holding neurons at 0 mV.

Data were collected using a Multiclamp 700B amplifier and Digidata 1440A digitizer (Axon Instruments) with pClamp 10 (Molecular Devices). Recordings were sampled at 10 kHz and filtered at 3 kHz for evoked recordings and 1kHz for spontaneous recordings. Series resistance and input resistance were monitored throughout the experiment by measuring the capacitive transient and steady-state deflection in response to a 5-mV test pulse, respectively. Series resistance was <25 MΩ, did not change more than 20% throughout a session and was not compensated. Data were analyzed in Python v3.7. Analysis was based on the average of ten sweeps. Currents were analyzed relative to the baseline holding current. EPSCs and IPSCs were quantified by measuring the peak response when cells were voltage clamped at –70 mV and 0 mV, respectively.

#### *In situ* hybridization

Mice were intracardially perfused with 20mL phosphate-buffered saline (PBS, Invitrogen) followed by 4% paraformaldehyde (PFA; Electron Microscopy Sciences). Brains were then extracted and placed in PFA for 24 hours and then sunk in a 30% sucrose solution for 3 days before being embedded in Optimal Cutting Temperature (OCT) compound and stored at -80℃. 20 µm-thick slices were prepared using a cryostat (Leica Microsystems) and mounted on Superfrost Plus microscope slides (Fisher Scientific). For *Vglut1* and *Vgat* labeling we processed the slices using RNAscope Multiplex Fluorescent Detection kit v2 (ACD Bio #323110) with the probe for *Vglut1* in C3 (ACD Bio #501101-C3) and *Vgat* in C2 (ACD Bio # 319191-C2) and TSA Vivid Dye 570 and 620, respectively. Retrogradely labelled vmPFC^CRH+^→BLA cells were stained using immunohistochemistry following the RNAscope protocol. Hydrophobic pen was used to create a barrier around brain slices before washing in 3x 10min washes in PBS, permeabilized for 2h in PBS with 0.5% Triton-X100 and incubated overnight at 4℃ using chicken anti-GFP primary antibody (1:2000; Aves Lab). The following day slices were washed in PBS and incubated for 2 hours in donkey anti-chicken Alexafluor 488 (1:1000; JacksonImmunoResearch) at RT. Following 30s incubation in DAPI slides were coverslipped and imaged on a Leica STELLARIS confocal microscope (Leica Microsystems) at 20x.

To verify successful knockdown in *Crh*-shRNA versus scrambled controls we processed the slices using the RNAscope Multiplex Fluorescent Detection kit v2 (ACD Bio #323110) with the probe for *Crh* in C2 (ACD Bio #316091-C2) and TSA Vivid Dye 620. mCherry labeling was done following the RNAscope protocol using immunohistochemistry. Following three washes in PBS slices were permeabilized for 2h in PBS with 0.5% Triton-X100 (PBST) and incubated overnight at 4℃ with a rabbit anti-RFP primary antibody (1:2000; Rockland Immunochemicals) diluted in PBST. The following day slices were washed for 10min 3x in PBS and incubated in Cy3 donkey anti-rabbit secondary antibody (1:1000; JacksonImmuno Research) for 2h at room temperature. DAPI was applied for 30s prior to mounting using fluoromount . Images were acquired using a Leica STELLARIS confocal microscope at 20x. All analyses were done in ImageJ and *Vglut1* and *Vgat* overlap with AAVrg CRH^+^ cells and *Crh* expression levels within mCherry positive cells were quantified using ImageJ version 1.54f.

To quantify *Crh* expression in *Vglut1*+ and *Vglut1*-cells in the vmPFC we processed the slices using the RNAscope kit with the probe for *Crh* in C2 (ACD Bio #316091-C2) and *Vglut1* in C1 (ACD Bio #501101) with the TSA Vivid Dye 620 and 650, respectively. Images were acquired using a Leica STELLARIS confocal microscope at 20x. All analyses were done in ImageJ, specifically the number of *Crh* expressing cells were quantified, along with their expression levels - calculated per *Vglut1*+ and *Vglut1*-cells. Quantification occurred in images acquired at 20x. Example images were acquired at 63x.

### Histology

Mice were intracardially perfused with 20mL phosphate-buffered saline (PBS, Invitrogen) followed by 4% paraformaldehyde (PFA; Electron Microscopy Sciences). Brains were then extracted and placed in PFA for 24 hours and then sunk in a 30% sucrose solution for 3 days before being embedded in Optimal Cutting Temperature (OCT) compound and stored at -80℃. 60 µm-thick slices were prepared using a cryostat For all experiments slices were permeabilized for 2h in PBST and incubated overnight at 4°C with primary antibodies diluted in PBST. The following day slices were washed in PBS for 5 min 3x, incubated for 2h at room temperature with secondary antibodies diluted in PBST, incubated in DAPI diluted 1:4000 in PBS and mounted and coverslipped with fluoromount. Images were acquired using either Leica DM6 scanning Microscope or Leica STELLARIS confocal.

For Fig 2A, Fig 3D, S 3A, Fig S4A,B, D, Fig S5A-D, Fig S7B,C a chicken anti-GFP primary (1:2000; Aves Lab) and donkey anti-chicken Alexafluor 488 (1:1000; Jackson Immunoresearch) were used.

For Fig S 3C a rabbit anti-pCREB (1:500; Cell Signaling Technology) primary antibody and Cy3 donkey anti-rabbit secondary antibody (1:100; JacksonImmuno Research, #711-165-152) were used.

For Fig 3D, Fig S5A-D, Fig 5F,M a rabbit anti-Fos (1:1000; Synaptic Systems) primary antibody and donkey anti-rabbit Alexa-Fluor 647 secondary antibody (1:1000; JacksonImmuno Research, #711-605-152) were used.

For Fig 1M, 4C, 5A,H, S2A, B, Fig S6A-C, Fig S8A,L, Fig S10A,B a rabbit anti-RFP (1:2000; Rockland Immunochemicals) and Cy3 donkey anti-rabbit secondary antibody (1:1000, JacksonImmuno Research) were used.

## QUANTIFICATION AND STATISTICAL ANALYSIS

Male and female mice were randomly assigned to experimental groups, ensuring both sexes were included in every group. For manipulation, experimental conditions (for example, control versus opsin) were divided among littermates to establish age-matched, litter-matched controls. Data collection and behavioral video and image analysis were performed by experimenters who were blind to the experimental groups. Behavioral data were analyzed blind to experimental conditions using automated analysis pipelines. For manual image analyses, researchers were blind to experimental manipulation.

All statistical tests were performed in GraphPad Prism (v10.3.1) or MATLAB (vR2022a, Mathworks). Our sample sizes are similar to those reported in previous publications; however, they were not predetermined with statistical methods. Summary graphs represent mean ± s.e.m. Shapiro-Wilk tests were used to determine normality, and nonparametric tests were used if found to be not normal. The statistical tests and values are reported in the legend of each figure. Paired t-tests were two-tailed, and significance was considered.

### Behavioral analysis

High resolution videos of PMA were collected at 50 Hz using Chameleon3 USB cameras (Teledyne FLIR) point tracking of videos was performed in DeepLabCut v2.0^77^, and behavior was analyzed using BehaviorDEPOT v1.3b^78^. Custom MATLAB code was used to quantify time on platform, latency to enter the platform, freezing, and locomotor speed. Open field, elevated plus maze, novel odor assays and RTPA were filmed using a webcam. Novel odor was recorded in real time by the experimenter. Open field, elevated plus maze and RTPA were tracked with automated video-tracking software, BioViewer.

#### Fiber photometry recordings during PMA

Animals were habituated to the optical tether 1 day before recordings. During PMA training and retrieval, we simultaneously imaged GCaMP6s and control fluorescence in the BLA using a commercial fiber photometry system and companion Synapse software controlling an RZ10x lock-in amplifier (Tucker David Technologies). Two excitation wavelengths (465 and 405 nm) were modulated at 211 and 566 Hz, filtered and combined by a fluorescence minicube (Doric Lenses). The combined excitation light was delivered via a 400µm core, 0.37 NA low-fluorescence was collected through the minicube and focused onto a femotwatt photoreceiver (Newport, Model 2151, gain set to DC LOW). LED power was such that 8- and 20 units of light were received by the system for the 465nm and 405nm channels, respectively. Fluorescence was sampled at 1,017 Hz and demodulated by the processor. Time stamps for experiment start and finish and each tone onset were collected using transistor-transistor logic (TTL) pulses sent from a custom MATLAB experiment designer (MathWorks). Signals were saved using Synapse software and exported to MATLAB for analysis.

#### Fiber photometry analysis

Data were preprocessed using a custom-written pipeline in MATLAB. Before analysis, signal was downsampled by 10x. Using the polyfit function, the isosbestic signal was fit to the 405nm signal, and this curve was subtracted from the 465nm channel. To align fiber photometry and behavioral data, a lookup table was generated using linear interpolation between each TTL pulse to identify which behavior frames line up with each photometry frame. Z-scores were calculated using a baseline period of -5 to 0s relative to tone onset. The average of all traces for an individual animal was calculated and used for analysis. To generate plots, each average trace was smoothed by averaging values from every 0.5s, and the mean ± s.e.m. of smoothed traces across animals were displayed. For shock response the area under the curve (AUC) and peak z-score were quantified during 0-15s and 0-3s after onset, respectively. The decay rate was measured by fitting an exponential curve from the peak shock response, decay is quantified as 1/τ.

#### Optogenetic manipulation during PMA

Before all experiments, mice were habituated to an optic tether (200µm core, 0.22 NA, Doric Lenses). No optogenetic manipulation occurred during PMA retrieval or the baseline three tones of training. For inhibition experiments vmPFC^CRH+^ projections to the BLA were photoinhibited with a blue laser (473 nm, constant; SLOC Lasers) controlled by BehaviorDEPOT fear conditioning experimenter MATLAB app during the shock period and 13s after the shock terminated, 15s total. This was chosen based on the bulk photometry results indicating that LBN mice displayed heightened activation in the BLA in response to the shock during this period. The light power delivered, as measured through an optic fiber preimplant, was set to an output of 10 mW of light. For stimulation experiments vmPFC^CRH+^ projections to the BLA were photoexcited with a blue laser (473 nm, 15 Hz, 50ms pulse width; SLOC Lasers) with the same light power.

#### Optogenetic manipulation during open field

On the day following PMA retrieval, animals were habituated for 10 minutes in a separate room. After connection to the blue laser (inhibition: 473 nm; constant; stimulation: 473 nm; 15 Hz), mice were placed in an open-field arena. Optogenetic manipulation began after a 1-min baseline, followed by eight alternating 30s laser on/off epochs, and concluded with a 60-s no-stimulation period, for a total session duration of 10 minutes.

### Whole-brain Analysis

#### DeepCOUNT Analysis Pipeline

DeepCOUNT^79^ was used to align and quantify cell density across the brain. Briefly, whole-brain image stacks from the 640nm channel from each brain were segmented in TrailMap using an Ai14-trained model. Images were registered to the template brain using elastix and transformix using the 488nm autofluorescence channel. MATLAB (Mathworks) was used to identify 3D maxima of the transformed probability map. Connected component analysis was used to reduce any maxima that consisted of multiple pixels into a single pixel per cell.

#### Cell Quantification

Regional CRH^+^ cell density was quantified in MATLAB by counting the number of labelled pixels (i.e. cells) in each brain region, then dividing this pixel count by the total number of pixels in the region. Regions were defined by a collapsed version of the LSFM atlas in which maximum granularity was balanced with the need to account for slight differences in registration which would lead to inaccurate quantification of small brain regions. This atlas was cropped on the anterior and posterior ends to match the amount of tissue visible in our data. Fiber tracts, ventricular systems, cerebellum, and olfactory bulb were excluded from analysis. For statistical analysis we focused on regions known to project to the BLA and contain CRH+ cell populations. Two-way repeated measures ANOVA with post-hoc multiple comparisons correction were performed to identify significant differences between brain regions.

#### Optogenetic manipulation during real-time place preference

To determine if optogenetic manipulation of vmPFC^CRH+^ projections impacted behavior beyond PMA, RTPP tests were performed the day following PMA retrieval for all optogenetic experiments. Following connection to the blue (inhibition: 473 nm; constant; stimulation: 473nm; 15 Hz), the mice were spaced in a place preference chamber (68 cm x 23 cm) for 20 min. For the first 10 min baseline period, mice were allowed to freely explore the apparatus and BioViewer software was used to track movement and determine which half of the chamber they preferred. This was used to determine which half of the chamber would receive laser stimulation or inhibition. For the following 10 min laser light was delivered on the preferred side of the chamber (stimulation) or the non-preferred side (inhibition). Results were calculated as percent change from baseline (test-baseline/baseline x 100).

#### Code and Data availability

All data reported in this paper will be shared by the lead contact upon request. BehaviorDEPOT used to analyze behavioral data is available at https://github.com/DeNardoLab/BehaviorDEPOT. DeepCOUNT used to analyze whole brain cell density is available at https://github.com/DeNardoLab/DeepTraCE. All other original code has been deposited at Github and is publicly available as of the date of publication. Any additional information required to reanalyze the data reported in this paper is available from the lead contact upon request.

## SUPPLEMENTAL INFORMATION

Extended Data Figures 1-10 and Extended Data Table 1.

## Notes

### Competing Interest Statement

The authors have declared no competing interest.

### Summary of Updates

This manuscript has been revised with additional experiments. These include Crh expression analysis in prefrontal cell types, Crh knockdown in glutamatergic prefrontal neurons, and pathway inhibition and Crh knockdown in adult mice.

