## Extended Data for "Early adversity promotes adolescent avoidance behavior by enhancing prefrontal–amygdala communication through CRH⁺ glutamatergic neurons"

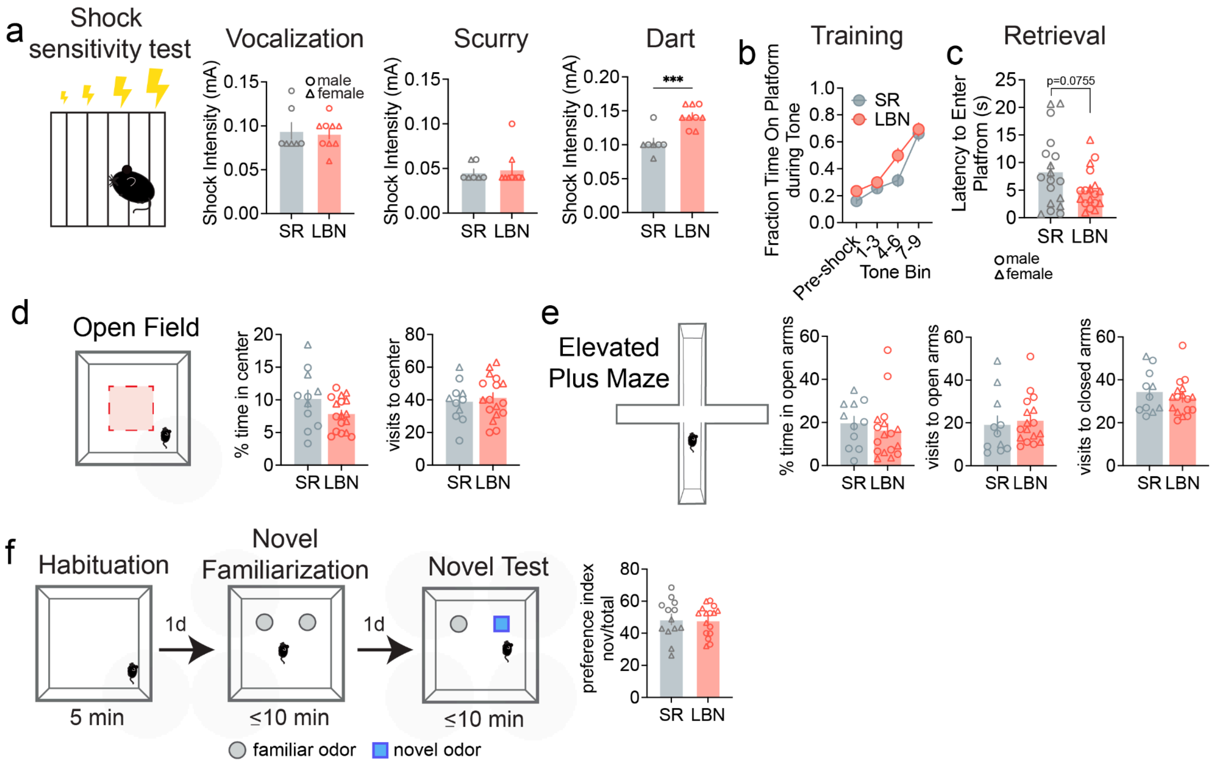


**Extended Data Figure 1. Effects of LBN on shock sensitivity, open field, elevated plus maze and novel odor assay in adolescent mice, Related to Figure 1.** (**a**) Summary data of shock levels required to elicit behavioral responses to progressive increases in shock intensity (SR, n=7; LBN, n=9 mice) (unpaired t-tests Vocalization $F_{6,8}$=2.02, p=0.770; Scurry: $F_{6,8}$=4.32, p=0.710; Dart: $F_{6,8}$=1.325, P=0.0004). (**b**) Fraction of time spent on the platform during PMA training (two-way repeated measures ANOVA with Sidak's post hoc test: trial effect $F_{2.48,86.71}$ = 69.69, P<0.0001; rearing effect $F_{1,35}$ = 4.79, P=0.036; interaction $F_{2.48,86.71}$= 1.92, P = 0.143). (**c**) Latency to enter the platform after tone onset in seconds (Mann-Whitney unpaired t-test $F_{17,17}$=3.38, p=0.0755). (**d**) Summary data of open field assay (SR, n=11; LBN, n=17 mice) including the time in center (unpaired t-test with Welch's correction $F_{10,16}$=3.110, p=0.1409), and visits to center (unpaired t-test $F_{16,10}$=1.146, p=0.637). (**e**) Summary data of elevated plus maze assay (SR, n=11; LBN, n=17) including time in the open arms (unpaired t-test $F_{16,10}$=1.579, p=0.487, visits to the open arms (unpaired t-test $F_{10,16}$=1.64, p=0.690) and visits to the closed arms (unpaired t-test $F_{10,16}$=1.373, p=0.450). (**f**) Summary data for novel odor assay (SR male, n=6; SR female, n=7, LBN male, n=8, LBN female, n=8 mice) measured as preference index (novel/total) (unpaired t-test $F_{12,15}$=1.74, p=0.889). Data are shown as mean +/- s.e.m. Triangles denote females, and circles denote males.

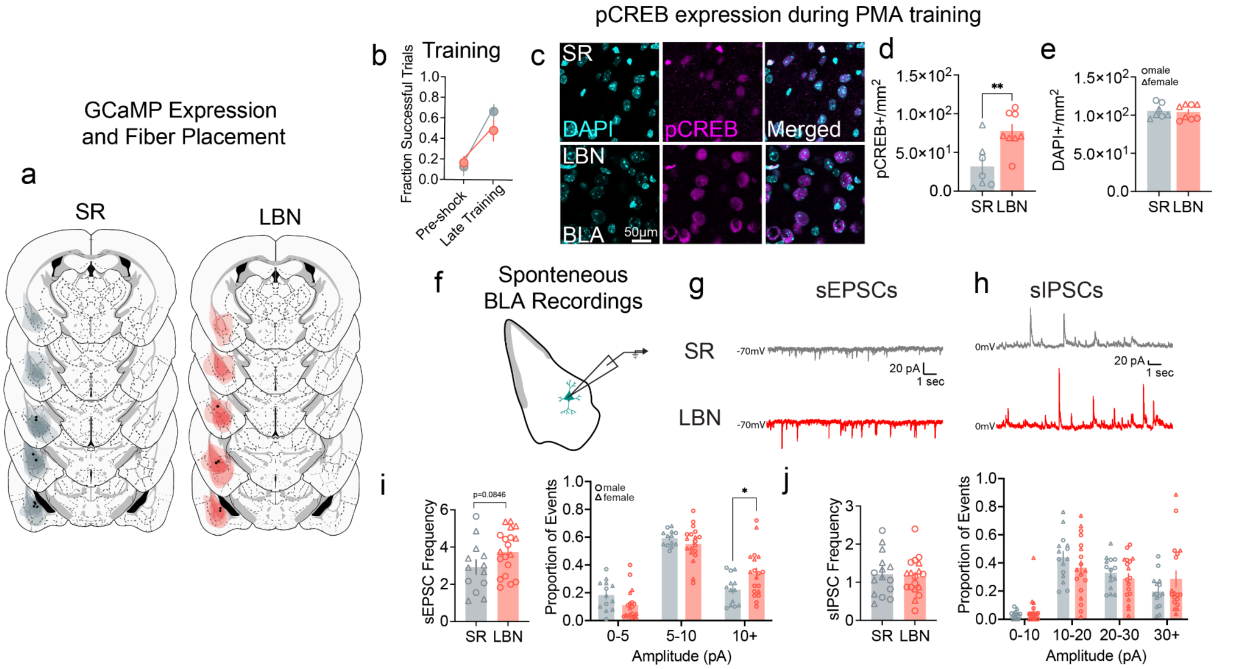


**Extended Data Figure 2. LBN enhances excitability via pCREB expression and spontaneous vesicle release in the BLA, Related to Figure 1.** (**a**) AAV-GCaMP6s viral expression and fiber implant placement in BLA for SR and LBN mice. (**b**) Fraction of successful trials during PMA training of animals taken for pCREB expression (SR, n=7; LBN, n=8 mice) (2-way ANOVA: trial effect $F_{1,29}$=25.25, p<0.0001; rearing effect $F_{1,29}$=0.71, p=0.407; interaction effect $F_{1,29}$=1.78, p=0.194). (**c**) Example of DAPI, pCREB and merged images taken at 63x in BLA following PMA training. (**d**) number of pCREB+ cells (unpaired t-test $F_{6,7}$=1.44, p=0.006) and (**e**) DAPI cells (unpaired t-test $F_{7,6}$=1.14, p=0.794) recorded per $mm^2$. (**f**) Schematic of whole cell patch clamp recordings of spontaneous postsynaptic currents. Representative traces of (**g**) sEPSCs and (**h**) sIPSCs in LBN and SR adolescent mice (EPSCs: SR, n=7 mice/13 cells; LBN, n=10 mice/19 cells; IPSCs: SR, n=7 mice/15 cells; LBN, n=10 mice/18 cells). Quantification of (**i**) sEPSC and (**j**) sIPSC frequency (sEPSC: unpaired t-test $F_{12,18}$=1.31, p=0.085; sIPSC: unpaired t-test $F_{14,17}$=1.32, p=0.737) and amplitudes (sEPSC: two-way ANOVA with Sidak's post hoc test: bin effect $F_{2,60}$=65.63, p<0.0001, rearing effect $F_{1,30}$=5.12, p=0.031, interaction effect $F_{2,60}$=3.61, p=0.033; sIPSC: two-way ANOVA with Sidak's post hoc test: bin effect $F_{3,93}$=22.14, p<0.0001, rearing effect $F_{1,31}$=0.087, p=0.770, interaction effect $F_{3,93}$=1,25, p=0.297). Data are shown as mean +/- s.e.m.; *P≤0.05, **P≤0.01. Triangles denote females, and circles denote males.

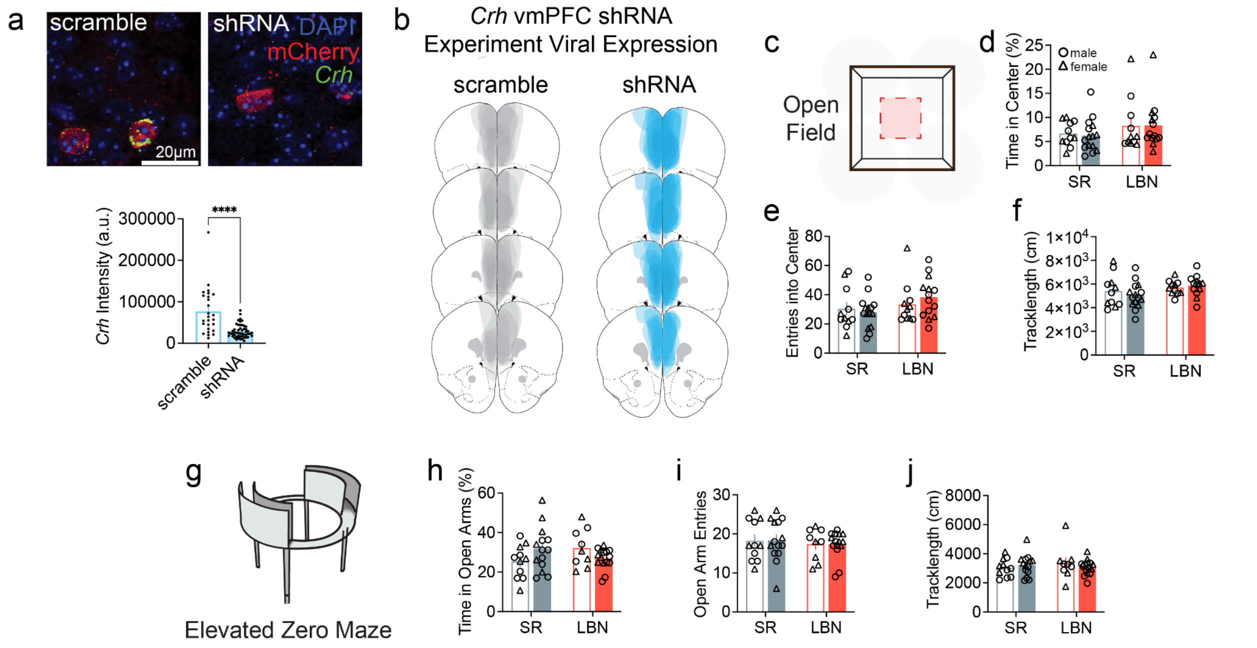


**Extended Data Figure 3. Histology and anxiety-like behaviors for *Crh* shRNA knockdown in vmPFC experiment, Related to Figure 2.** (**a**) Example images of in situ hybridization labelling Crh expression (green) in mCherry+ cells (red) in scramble controls versus shRNA knockdown mice. Quantification of Crh expression between conditions (Mann-Whitney test (two-tailed) p=0.0007; scramble, n=26 cells/2 mice; shRNA, n=51 cells/4 mice). (**b**) AAV-shRNA viral expression in mPFC of CRH-cre adolescent mice. (**c**) Summary data of open field assay (SR scramble, n=11; SR shRNA, n=15, LBN scramble, n=11; LBN shRNA, n=14 mice) including the (**d**) time in center (two-way ANOVA: interaction effect $F_{1,47}$=0.036, p=0.851, rearing effect $F_{1,47}$=2.36, p=0.133, knockdown effect $F_{1,47}$=0.013, p=0.910), (**e**) entries into the center (two-way ANOVA: interaction effect $F_{1,47}$=0.82, p=0.271, rearing effect $F_{1,47}$=3.06, p=0.087, knockdown effect $F_{1,47}$=0.11, p=0.737) and (**f**) tracklength in cm (two-way ANOVA: interaction effect $F_{1,47}$=0.85, p=0.362, rearing effect $F_{1,47}$=3.65 p=0.062, knockdown effect $F_{1,47}$=0.11, p=0.740). (**g**) Summary data of elevated plus maze assay (SR scramble, n=11; SR shRNA, n=14, LBN scramble, n=9; LBN shRNA, n=15 mice) including (**h**) time in the open arms (two-way ANOVA: interaction effect $F_{1,45}$=5.11, p=0.023, rearing effect $F_{1,45}$=0.135 p=0.715, rearing effect $F_{1,45}$=0.057, p=0.812), (**i**) visits to the open arms (two-way ANOVA: interaction effect $F_{1,45}$=0.0043, p=0.948, rearing effect $F_{1,45}$=0.49 p=0.488, knockdown effect $F_{1,45}$=0.0043, p=0.948) and (**j**) tracklength (two-way ANOVA: interaction effect $F_{1,45}$=1.0, p=0.323, rearing effect $F_{1,45}$=0.144 p=0.701, knockdown effect $F_{1,45}$=0.045 p=0.832. Data are shown as mean +/- s.e.m. Triangles denote females, and circles denote males.

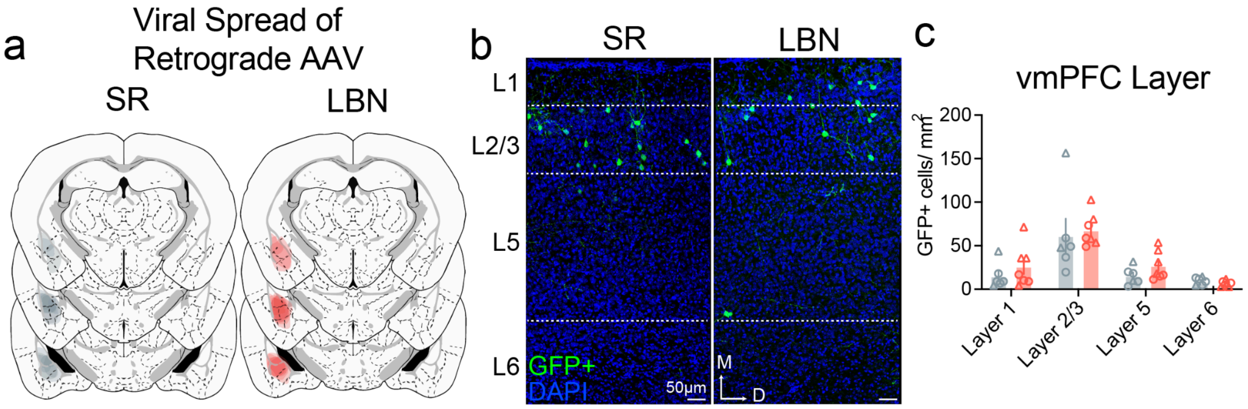


**Extended Data Figure 4. Distribution of CRH+ cells that project to the BLA in throughout the brain and within the vmPFC, Related to Figure 3.** (**a**) AAVrg-FLEX-GFP viral expression in BLA for SR and LBN animals. (**b**) Example images of CRH+ that project to the BLA in layers of the vmPFC (SR, n=6; LBN, n=7 mice). (**c**) Quantification of layer analysis (two-way repeated measures ANOVA with Sidak's post hoc test: layer effect $F_{1.48,16.31}$ = 19.41, P=0.001; rearing effect $F_{1,11}$ = 0.99, P=0.342; interaction $F_{1.48,16.31}$ = 0.35, P = 0.647). All data points represent biological replicates. Data are shown as mean +/- s.e.m. Triangles denote females, and circles denote males.

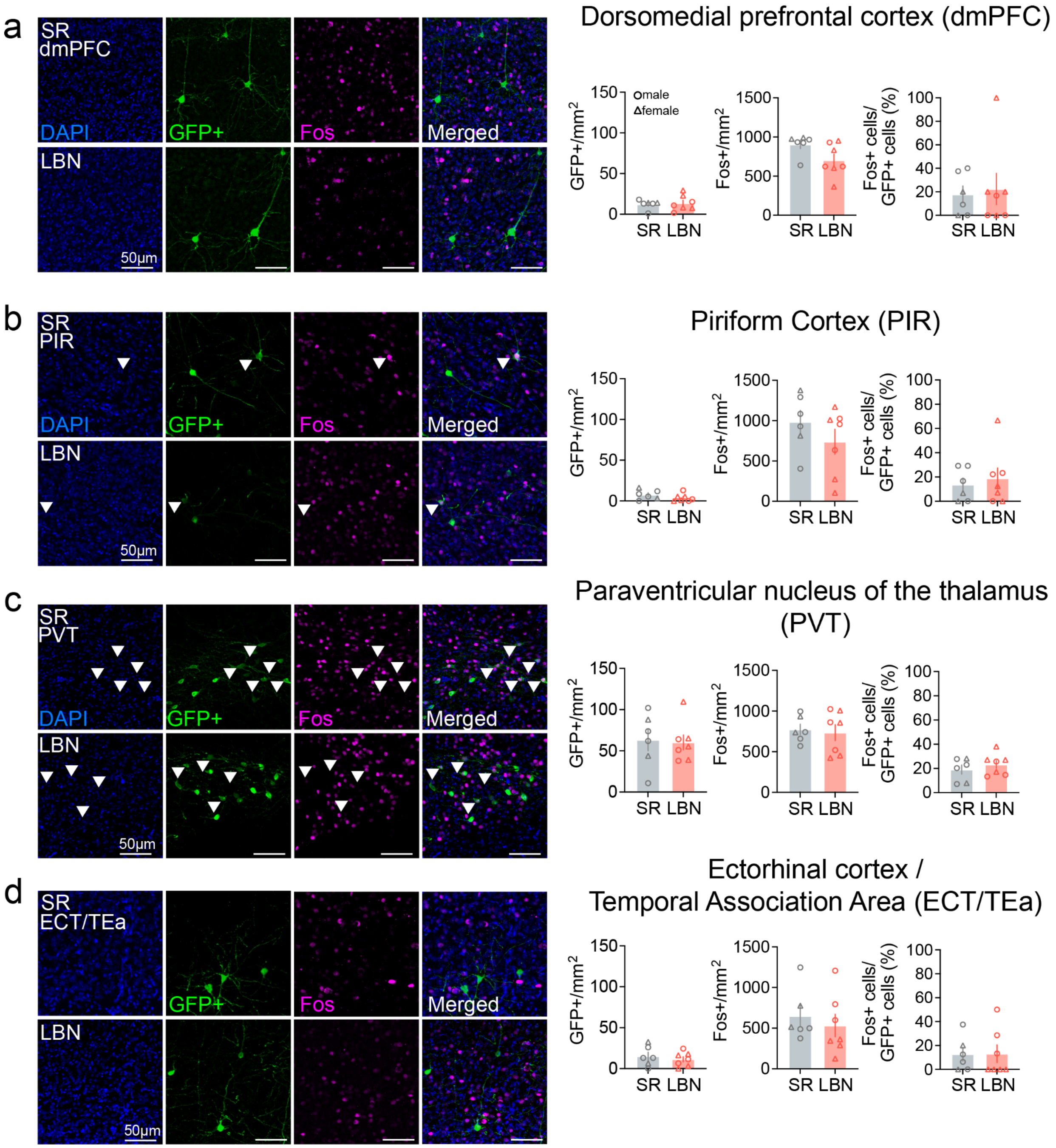


**Extended Data Figure 5. Activation of retrogradely labelled regions following PMA training. Related to Figure 3.** (**a**) Examples of DAPI (blue), retrogradely labelled CRH$^+$ cells (green), Fos+ cells (magenta) and merged images in the dmPFC. Quantification of the number of GFP$^+$ cells (unpaired t-test $F_{5,6}$=1.64, p=0.231), Fos+ cells (unpaired t-test $F_{6,5}$=2.52, p=0.070) and the percent of GFP$^+$/Fos+ cells (unpaired t-test $F_{6,5}$=2.96, p=0.779). (**b**) Examples of DAPI (blue), retrogradely labelled CRH$^+$ cells (green), Fos+ cells (magenta) and merged images in the piriform cortex (PIR). Quantification of the number of GFP$^+$ cells (unpaired t-test $F_{6,5}$=2.78, p=0.756), Fos+ cells (unpaired t-test $F_{6,5}$=1.35, p=0.280) and the percent of GFP$^+$/Fos+ cells (unpaired t-test $F_{6,5}$=2.93, p=0.637). (**c**) Examples of DAPI (blue), retrogradely labelled CRH$^+$ cells (green),

Fos+ cells (magenta) and merged images in the paraventricular nucleus of the thalamus (PVT). Quantification of the number of GFP$^+$ cells (unpaired t-test $F_{5,6}$=1.82, p=0.849), Fos+ cells (unpaired t-test $F_{6,5}$=2.61, p=0.762) and the percent of GFP$^+$/Fos+ cells (unpaired t-test $F_{5,6}$=1.14, p=0.418). (**d**) Examples of DAPI (blue), retrogradely labelled CRH$^+$ cells (green), Fos+ cells (magenta) and merged images in the ectorhinal cortex and temporal association area (ECT/TEa). Quantification of the number of GFP$^+$ cells (unpaired t-test $F_{5,6}$=1.94, p=0.525), Fos+ cells (unpaired t-test $F_{6,5}$=1.32, p=0.554) and the percent of GFP$^+$/Fos+ cells (unpaired t-test $F_{6,5}$=1.85, p=0.967). White arrows point to GFP+/Fos+ cells. All data points represent biological replicates. Data are shown as mean +/- s.e.m. Triangles denote females, and circles denote males.

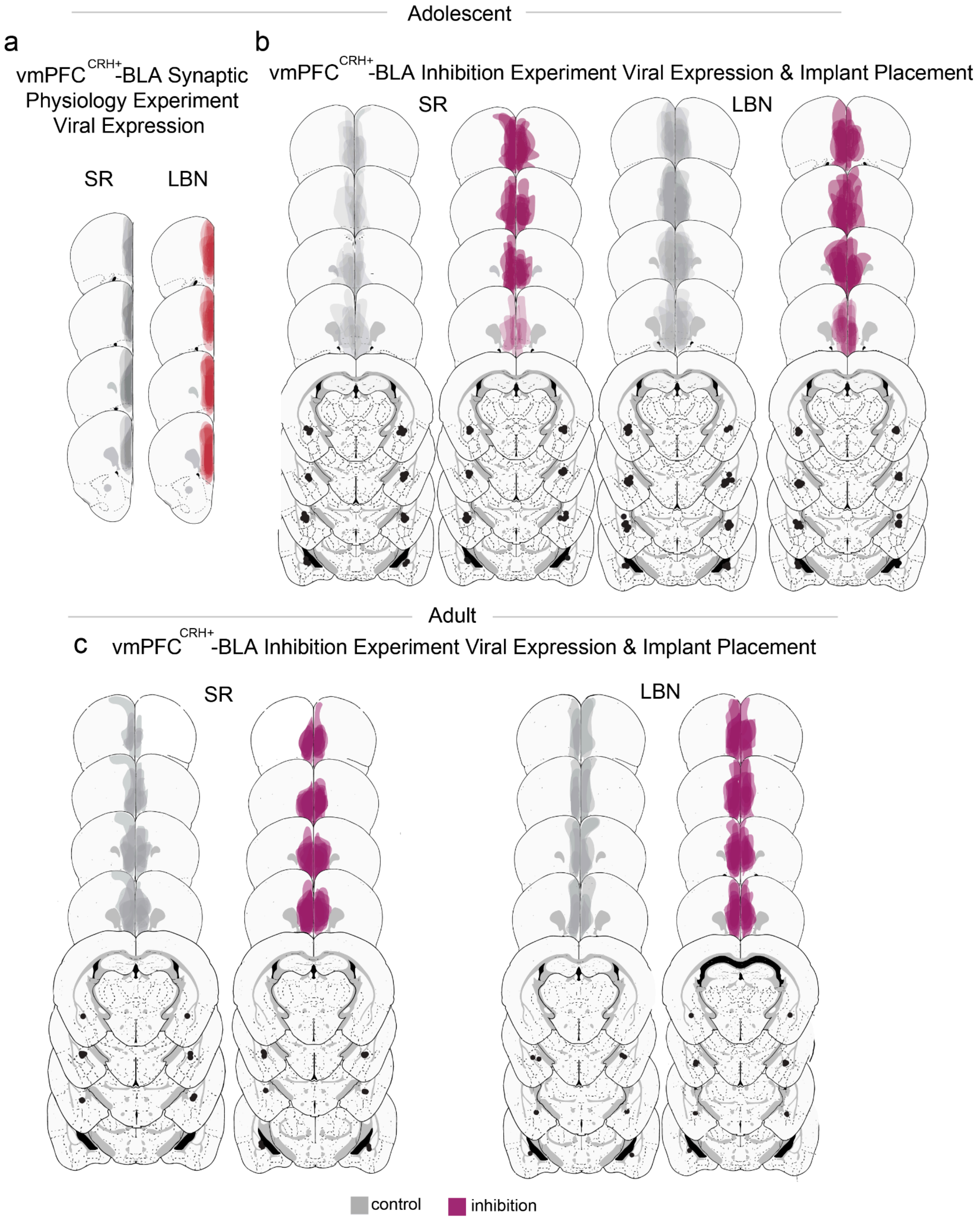


**Extended Data Figure 6. Histology for vmPFC$^{CRH+}$ → BLA synaptic physiology and optogenetic manipulation experiments, Related to Figures 3 and 4.** (**a**) AAV-ChR2 viral expression in mPFC of SR and LBN mice for synaptic physiology experiments. (**b**) AAV-PdCO or control viral expression in mPFC of SR and LBN adolescent mice and fiber implant placement above BLA. (**c**) AAV-PdCO or control viral expression in mPFC of SR and LBN adult mice and fiber implant placement above BLA.

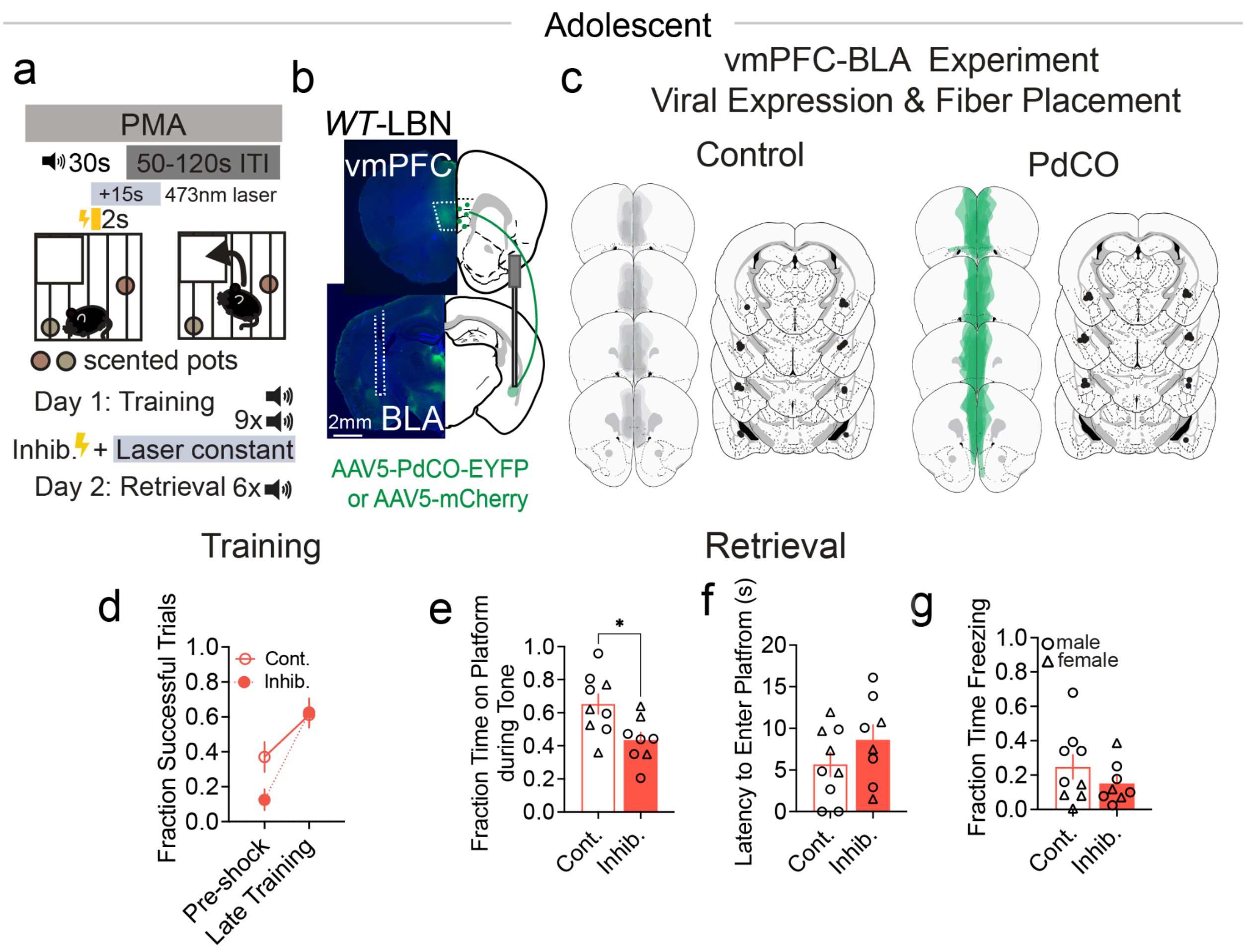


**Extended Data Figure 7. Inhibition of vmPFC–BLA pathway reduces threat avoidance in LBN mice, Related to Figure 4.** (**a**) Schematic representation of PMA paradigm with constant laser stimulation during the shock period and 13 seconds post shock during training. (**b**) Schematic and example images of AAV-PdCO expression and bilateral fiber optic implants in the BLA of wild-type LBN adolescent mice (Cont., n=9; Inhib., n=8). (**c**) AAV-PdCO viral expression in vmPFC in LBN mice and fiber implant placement above BLA. (**d**) Fraction of successful trials during PMA training averaged for the pre-shock baseline and the last four tones of the session (two-way ANOVA: trial effect $F_{1,15}$=41.52, p<0.0001; opsin effect $F_{1,15}$=1.55, p=0.232; interaction effect $F_{1,15}$=5.09, p=0.040). (**e-g**), Behavioral performance during PMA retrieval. (**e**) Fraction of time on the platform averaged across the session (unpaired t-test $F_{8,7}$=3.67, p=0.836). (**f**) Latency to enter the platform following tone onset (unpaired t-test $F_{7,8}$=1.36, p=0.213). (**g**) Fraction of time freezing during the tone (unpaired t-test $7_{8,7}$=3.18, p=0.270). Data are shown as mean +/- s.e.m.; *P<0.05. All data points represent biological replicates. Data are shown as mean +/- s.e.m. Triangles denote females, and circles denote males.

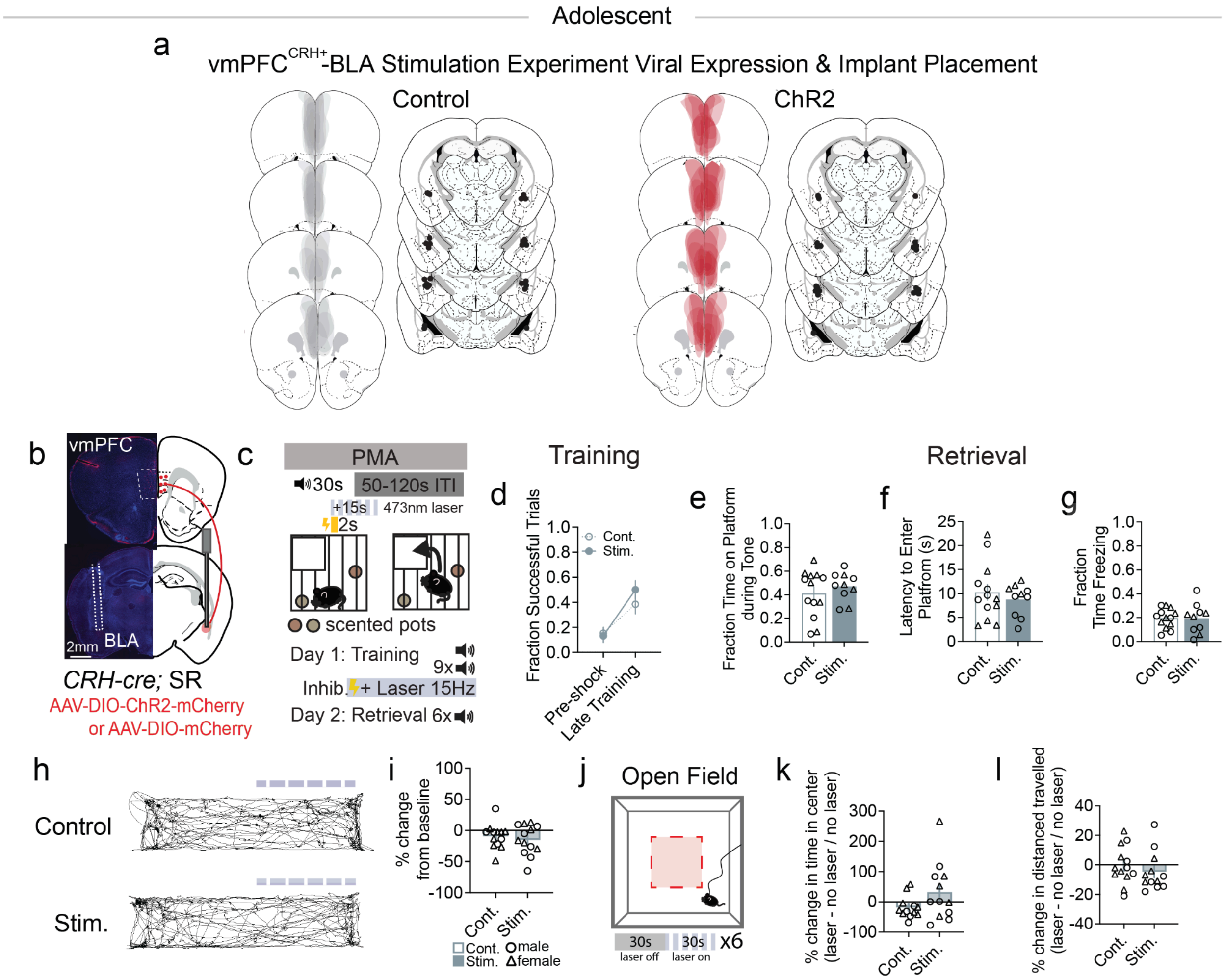


**Extended Data Figure 8. Stimulation of the vmPFC$^{CRH+}$→ BLA pathway is not sufficient to drive increased avoidance in SR mice, Related to Figure 4.** (**a**) AAV-ChR2 viral expression in mPFC of SR and LBN mice for stimulation experiments. (**b**) Schematic and example images of AAV-DIO-ChR2 expression and bilateral fiber optic implants in the BLA of CRH-cre mice. (**c**) Schematic representation of PMA with 15Hz laser stimulation during the shock period and 13 seconds post shock during training (Cont., n=13; Stim., n=10 mice). (**d**) Fraction of successful trials during PMA training averaged for the pre-shock baseline and the last four tones, only in SR mice (two-way ANOVA: trial effect $F_{1,21}$=23.77, p<0.0001; opsin effect $F_{1,21}$=0.45, p=0.510; interaction effect $F_{1,21}$=1.23, p=0.280). (**e-g**), Behavioral performance during PMA retrieval. (**e**) Fraction of time on the platform averaged across the session (unpaired t-test $F_{12,9}$=2.73, p=0.426). (**f**) Latency to enter the platform following tone onset (unpaired t-test $F_{12,9}$=3.18, p=0.481). (**g**) Fraction of time freezing during the tone (unpaired t-test $F_{12,9}$=2.33, p=0.976). (**h**) Representative mouse trajectory maps during RTPA with vmPFC$^{CRH+}$→BLA stimulation for control and ChR2-expressing mice in SR (cont., n=13; stim., n=13 mice). (**i**) Summary data of fraction of time spent on the laser-paired side of the chamber for stimulation experiments (unpaired t-test with Welch's correction: $F_{12,12}$=5.91, p=0.933). (**j**) Diagram of OF with 30 second 15Hz laser stimulation interleaved for 10 min (cont., n=13; stim., n=12 mice). Summary data of OF quantified as (**k**) percent change in time in center (unpaired t-test with Welch's correction: $F_{11,11}$=6.84, p=0.111) and (**l**) percent distance travelled as a function of laser stimulation (unpaired t-test: $F_{11,12}$=1.07, p=0.405). All data points represent biological replicates. Data are shown as mean +/- s.e.m.; Triangles denote females, and circles denote males.

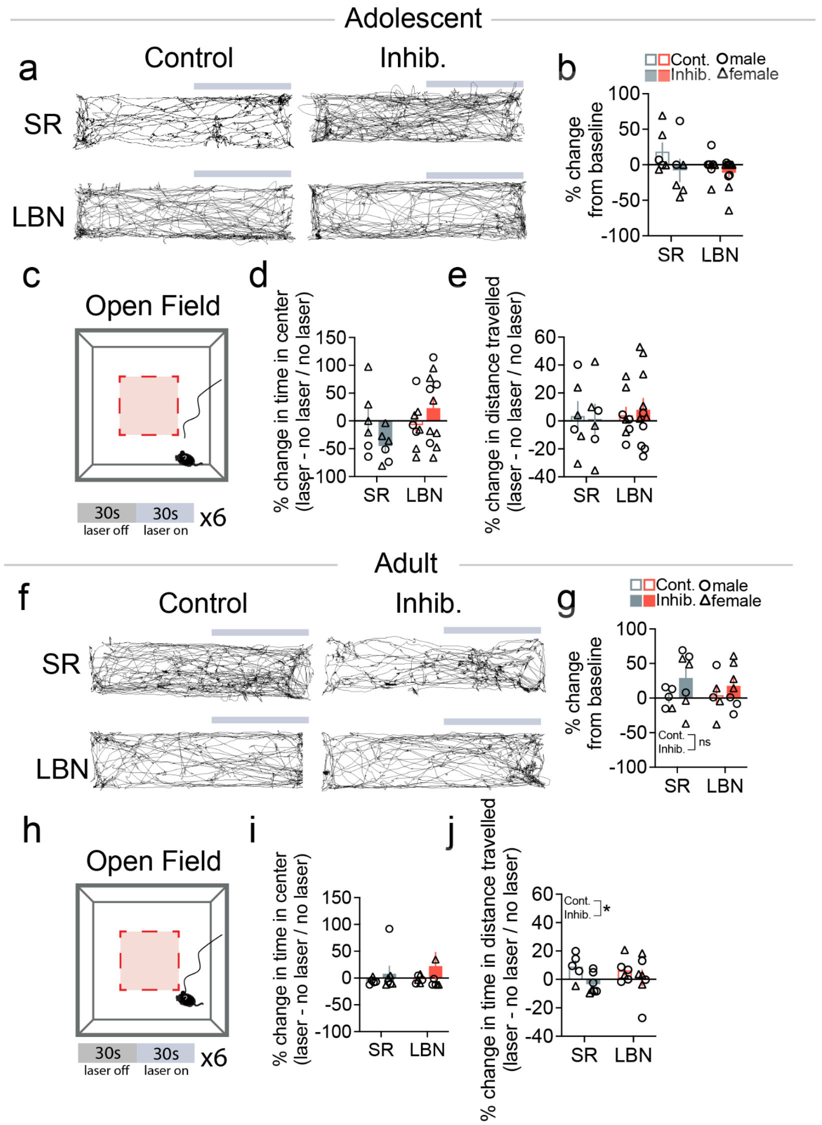


**Extended Data Figure 9. Inhibition of vmPFC$^{CRH+}$→ BLA drives Real Time Place Preference only in adults, Related to Figure 4.** (**a**) Representative mouse trajectory maps during RTPP in adolescents with vmPFC$^{CRH+}$→BLA inhibition for control and PdCO-expressing mice in SR and LBN groups (SR cont., n=6; SR inhib., n=6; LBN cont., n=8, LBN inhib., n=11 mice). (**b**) Summary data of fraction of time spent on the side of the laser-paired side of the chamber for inhibition experiments (two-way ANOVA: rearing effect $F_{1,27}$=1.43, p=0.242; opsin effect $F_{1,27}$=3.49, p=0.073; interaction effect $F_{1,27}$=0.391, p=0.761). (**c**) Diagram of OF with 30 second constant laser stimulation interleaved for 10 min in adolescents. Summary data of OF quantified as (**d**) percent change in time in center (two-way ANOVA: rearing effect $F_{1,27}$=2.50, p=0.126; opsin effect $F_{1,27}$=0.141, p=0.711; interaction effect $F_{1,27}$=3.79, p=0.062) and (**e**) percent distance travelled as a consequence of laser stimulation (two-way ANOVA: rearing effect $F_{1,27}$=0.166, p=0.687; opsin effect $F_{1,27}$=0.117, p=0.915; interaction effect $F_{1,27}$=0.122, p=0.730). (**f**) Representative mouse trajectory maps during RTPP in adults with vmPFC$^{CRH+}$→BLA inhibition for control and PdCO-expressing mice in SR and LBN groups (SR cont., n=5; SR inhib., n=7; LBN cont., n=5, LBN inhib., n=7 mice). (**g**) Summary data of fraction of time spent on the side of the laser-paired side of the chamber for inhibition experiments (two-way ANOVA: rearing effect $F_{1,20}$=0.078, p=0.883; opsin effect $F_{1,20}$=2.60, p=0.123; interaction effect $F_{1,20}$=0.033, p=0.572). (**h**) Diagram of OF with 30 second constant laser stimulation interleaved for 10 min in adults. Summary data of OF quantified as (**i**) percent change in time in center (two-way ANOVA: rearing effect $F_{1,21}$=0.265, p=0.612; opsin effect $F_{1,21}$=1.28, p=0.270; interaction effect $F_{1,21}$=0.103, p=0.751) and (**j**) percent distance travelled as a consequence of laser stimulation (two-way ANOVA: rearing effect $F_{1,21}$=0.037, p=0.849; opsin effect $F_{1,21}$=4.479, p=0.0.046; interaction effect $F_{1,21}$=0.725, p=0.404). All data points represent biological replicates. Data are shown as mean +/- s.e.m.; Triangles denote females, and circles denote males.

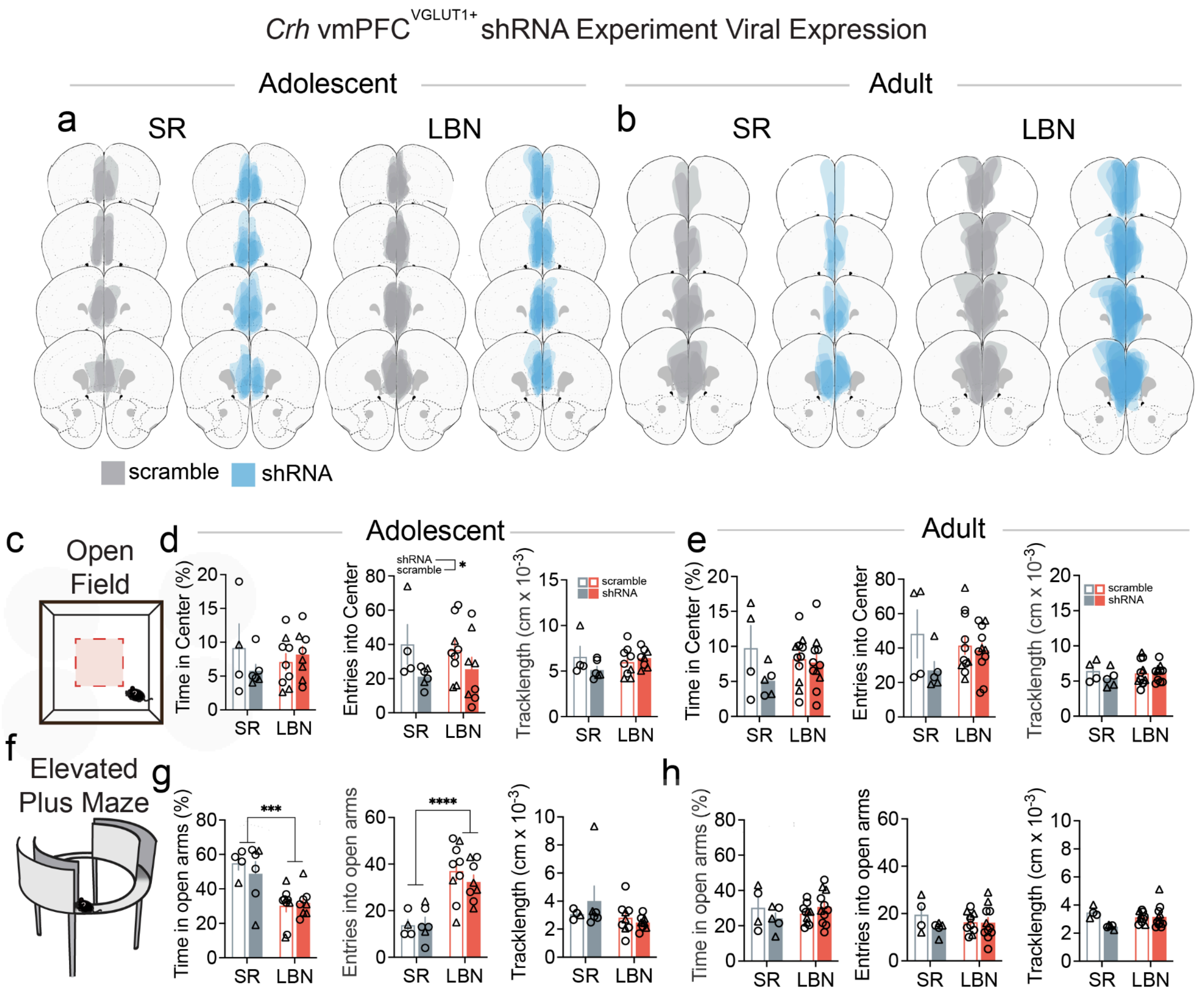


**Extended Data Figure 10. *Crh* knocdown in vmPFC$^{VGLUT1+}$ cells doesn't impact anxiety-like behavior, Related to Figure 5.** (**a**) AAV-shRNA viral expression in mPFC of VGLUT1-cre adolescent and (**b**) adult mice. (**c**) Schematic of OF in adolescent (SR cont., n=4; SR inhib., n=6; LBN cont., n=9, LBN inhib., n=8 mice) and adult (SR cont., n=4; SR inhib., n=5; LBN cont., n=11, LBN inhib., n=11 mice) mice. (**d**) Quantification of time in center (two-way ANOVA: knockdown effect $F_{1,23}$=0.494, p=0.489; rearing effect $F_{1,23}$=0.011, p=0.916; interaction effect $F_{1,23}$=1.86, p=0.186), entries into center (two-way ANOVA: knockdown effect $F_{1,23}$=4.65, p=0.0418; rearing effect $F_{1,23}$=0.0049, p=0.945; interaction effect $F_{1,23}$=0.30, p=0.588) and total tracklength travelled in cm in adolescent mice (two-way ANOVA: knockdown effect $F_{1,23}$=0.74, p=0.40; rearing effect $F_{1,23}$=0.409, p=0.529, interaction effect $F_{1,23}$=2.62, p=0.119) for vmPFC$^{VGLUT1}$+ Crh knockdown experiment. (**e**) Quantification of time in center (two-way ANOVA: knockdown effect $F_{1,16}$=1.82, p=0.106; rearing effect $F_{1,16}$=0.296, p=0.594; interaction effect $F_{1,16}$=1.52, p=0.235), entries into center (two-way ANOVA: knockdown effect $F_{1,16}$=1.71, p=0.210; rearing effect $F_{1,16}$=0.0004, p=0985; interaction effect $F_{1,16}$=1.63, p=0.221) and total tracklength travelled in cm in adolescent mice (two-way ANOVA: knockdown effect $F_{1,16}$=0.14, p=0.717; rearing effect $F_{1,16}$=0.078, p=0.784, interaction effect $F_{1,16}$=1.50, p=0.238) for vmPFC$^{VGLUT1}$+ Crh knockdown experiment. (**f**) Schematic of EZM. (**g**) Quantification of time in open arms (two-way ANOVA: knockdown effect $F_{1,23}$=0.226, p=0.639; rearing effect $F_{1,23}$=19.71, p=0.0002; interaction effect $F_{1,23}$=0.674, p=0.42), entries into open arms (two-way ANOVA: knockdown effect $F_{1,23}$=0.282, p=0.6004; rearing effect $F_{1,23}$=29.45, p<0.0001; interaction effect $F_{1,23}$=0.469, p=0.500) and total tracklength travelled (two-way ANOVA: knockdown effect $F_{1,23}$=0.351, p=0.559; rearing effect $F_{1,23}$=2.16, p=0.155; interaction effect $F_{1,23}$=1.43, p=0.244) for vmPFC$^{VGLUT1+}$ Crh knockdown experiment in adolescent mice. (**h**) Quantification of time in open arms (two-way ANOVA: knockdown effect $F_{1,27}$=2.74, p=0.109; rearing effect $F_{1,27}$=102, p=0.751; interaction effect $F_{1,27}$=1.78, p=0.193), entries into open arms (two-way ANOVA: knockdown effect $F_{1,27}$=3.15, p=0.087;

rearing effect $F_{1,27}$=0.12, p=0.723; interaction effect $F_{1,27}$=1.85, p=0.185) and total tracklength travelled (two-way ANOVA: knockdown effect $F_{1,27}$=0.72, p=0.402; rearing effect $F_{1,27}$=0.14, p=0.708; interaction effect $F_{1,27}$=0.89, p=0.354) for vmPFC$^{VGLUT1+}$ Crh knockdown experiment in adult mice. All data points represent biological replicates. Data are shown as mean +/- s.e.m.; Triangles denote females, and circles denote males.

**Extended Data Table 1, Related to Figure 2.**
**Adolescent CRH+ cells counts normalized to region volume for SR and LBN mice, Related to Figure 2.**

| Region | SR 1 | SR 3 | SR 4 | SR 5 | SR 6 | SR 8 | LBN 1 | LBN 2 | LBN 3 | LBN 4 | LBN 5 | LBN 6 | LBN 7 |
|---|---|---|---|---|---|---|---|---|---|---|---|---|---|
| Primary motor area' | 9240.23 | 7779.58 | 10729.99 | 7983.33 | 8737.47 | 9078.82 | 6543.84 | 9348.72 | 8840.67 | 9078.82 | 7051.90 | 8192.37 | 8562.83 |
| Secondary motor area' | 9125.39 | 7285.18 | 10451.59 | 8068.44 | 8119.62 | 8613.61 | 6125.87 | 9265.58 | 8811.65 | 9169.90 | 7423.14 | 8010.59 | 8825.00 |
| Primary somatosensory area' | 6766.05 | 6132.42 | 8139.13 | 6555.70 | 7308.06 | 7337.74 | 5525.89 | 7732.63 | 7407.42 | 7832.00 | 5969.82 | 6396.97 | 7002.21 |
| Supplemental somatosensory area' | 9917.49 | 8538.47 | 10503.39 | 8820.71 | 9224.41 | 9871.05 | 8995.76 | 9806.74 | 10528.40 | 10564.13 | 8216.94 | 8567.05 | 9163.68 |
| Gustatory areas' | 6906.89 | 5479.11 | 7585.08 | 6960.43 | 6817.65 | 7852.79 | 6746.26 | 6853.34 | 7263.83 | 7906.33 | 6068.06 | 5746.81 | 6121.61 |
| Visceral area' | 10744.57 | 9491.51 | 11738.37 | 9693.15 | 10917.40 | 9865.99 | 9693.15 | 11133.44 | 11032.62 | 10154.05 | 8440.10 | 9160.25 | 10082.03 |
| Dorsal auditory area' | 11125.93 | 11151.69 | 10894.14 | 11151.69 | 11254.70 | 10404.81 | 11254.70 | 11563.76 | 11718.28 | 13109.03 | 9271.61 | 10147.26 | 11692.53 |
| Primary auditory area' | 11250.78 | 9835.02 | 10542.90 | 10151.31 | 11130.29 | 10618.21 | 10663.39 | 10994.74 | 10844.13 | 11115.23 | 9654.29 | 9443.43 | 11190.54 |
| Posterior auditory area' | 11639.13 | 10254.74 | 12510.78 | 11536.58 | 11946.77 | 11690.40 | 10767.47 | 11331.48 | 10767.47 | 11792.95 | 9998.37 | 10459.83 | 11741.67 |
| Ventral auditory area' | 9166.62 | 8398.37 | 8607.89 | 8276.15 | 9026.94 | 9219.00 | 7717.42 | 9149.16 | 8276.15 | 9672.96 | 7699.96 | 7769.80 | 9009.48 |
| Anterolateral visual area' | 10941.49 | 11983.54 | 11542.67 | 10741.10 | 11382.36 | 11662.91 | 11702.99 | 11302.20 | 13466.45 | 11863.30 | 10139.92 | 11101.81 | 11983.54 |
| Anteromedial visual area' | 5788.88 | 6852.14 | 8309.20 | 5198.17 | 5119.41 | 6930.90 | 3268.55 | 6930.90 | 6064.54 | 6615.86 | 5670.74 | 6418.96 | 10160.07 |
| Lateral visual area' | 12582.47 | 12884.45 | 13563.91 | 11273.90 | 12381.16 | 13639.40 | 11525.55 | 12733.46 | 14016.88 | 13337.42 | 12079.18 | 11777.20 | 12808.96 |
| Primary visual area' | 13222.84 | 12197.94 | 14033.88 | 12218.11 | 13828.09 | 14025.81 | 11665.31 | 13037.23 | 13045.30 | 13436.69 | 11826.71 | 12452.14 | 13634.41 |
| Posterolateral visual area' | 12286.86 | 9702.02 | 10658.06 | 11684.91 | 10551.83 | 10905.92 | 10799.69 | 10233.15 | 10693.47 | 12145.23 | 10622.65 | 10091.52 | 11153.78 |
| posteromedial visual area' | 8729.89 | 8499.39 | 13109.23 | 7490.99 | 8528.21 | 8557.02 | 6338.53 | 7980.79 | 9507.80 | 8672.26 | 8096.03 | 8067.22 | 8701.07 |
| dmPFC' | 12624.01 | 9509.11 | 14461.75 | 12210.95 | 10874.90 | 12501.80 | 10697.82 | 14499.74 | 10942.24 | 11507.05 | 10293.21 | 11696.79 | 13037.06 |
| vmPFC' | 20549.44 | 11677.10 | 17636.95 | 15371.68 | 17097.66 | 14832.19 | 9681.37 | 13753.63 | 12701.79 | 14913.20 | 13753.63 | 12701.79 | 12108.44 |
| Orbital area, lateral part' | 6364.02 | 5071.33 | 7408.12 | 5170.77 | 5991.13 | 5916.55 | 4822.74 | 6202.44 | 6910.93 | 8240.91 | 4586.57 | 5058.90 | 5580.95 |
| Orbital area, medial part' | 5924.30 | 3748.02 | 6770.62 | 4014.01 | 5779.21 | 5102.15 | 4860.34 | 5295.60 | 6601.36 | 10857.18 | 4352.54 | 4932.88 | 5174.69 |
| Orbital area, ventrolateral part' | 4850.84 | 4338.15 | 7749.52 | 4752.25 | 6526.95 | 6428.35 | 4771.97 | 6310.04 | 9050.96 | 9977.75 | 4771.97 | 5955.10 | 5895.94 |
| Agranular insular area, dorsal part' | 7910.62 | 6567.75 | 9099.15 | 7115.70 | 7216.03 | 8188.46 | 6891.89 | 8443.14 | 8489.45 | 9284.37 | 6606.34 | 6961.35 | 7254.62 |
| Agranular insular area, | 9410.15 | 7914.95 | 10356.65 | 8902.61 | 9958.85 | 9286.69 | 8532.24 | 10000.00 | 10027.43 | 10507.54 | 8010.97 | 8230.45 | 9423.87 |

| | | | | | | | | | | | | | |
|---|---|---|---|---|---|---|---|---|---|---|---|---|---|
| posterior part' | | | | | | | | | | | | | |
| Agranular insular area, ventral part' | 9072.01 | 8442.01 | 10926.02 | 8964.01 | 9738.01 | 10926.02 | 8442.01 | 9828.01 | 10350.02 | 10350.02 | 7722.01 | 8856.01 | 9450.01 |
| Retrosplenial area' | 10740.43 | 10403.14 | 13110.68 | 10146.31 | 10192.73 | 11845.10 | 6857.04 | 11876.05 | 10028.73 | 10721.86 | 9750.24 | 11708.95 | 12640.35 |
| Rostrolateral visual area' | 5436.08 | 5198.59 | 5436.08 | 5172.20 | 5647.19 | 5568.03 | 4248.59 | 5911.08 | 6069.41 | 6386.08 | 4380.53 | 4301.37 | 5726.36 |
| Temporal association areas' | 8204.37 | 6845.36 | 6905.76 | 6956.10 | 7640.63 | 7348.70 | 7640.63 | 6976.23 | 7399.03 | 7248.03 | 6633.96 | 6784.96 | 7862.10 |
| Perirhinal area' | 7438.64 | 5406.62 | 6168.63 | 5842.05 | 5551.76 | 5805.77 | 5624.34 | 5551.76 | 5950.91 | 5588.05 | 5624.34 | 5188.90 | 6313.77 |
| Ectorhinal area' | 7811.94 | 7882.00 | 7846.97 | 7303.99 | 8284.86 | 7776.91 | 7391.57 | 8337.41 | 7444.11 | 7846.97 | 6708.46 | 7409.08 | 8670.20 |
| Accessory olfactory bulb' | 1394.55 | 1704.45 | 1756.10 | 1342.90 | 1033.00 | 2375.90 | 1549.50 | 2117.65 | 1394.55 | 7437.60 | 1652.80 | 1394.55 | 2427.55 |
| Anterior olfactory nucleus' | 6723.86 | 4827.39 | 8340.18 | 6048.60 | 6084.52 | 6228.19 | 4489.76 | 7176.43 | 6925.01 | 8835.85 | 4820.21 | 5653.51 | 5725.34 |
| Taenia tecta' | 15707.37 | 8155.75 | 11352.60 | 10421.24 | 13391.54 | 11503.64 | 9212.98 | 11075.71 | 11000.20 | 14750.83 | 9162.63 | 9313.67 | 8432.64 |
| Piriform area' | 9330.09 | 8361.02 | 10392.12 | 8397.64 | 9631.51 | 9448.41 | 8121.57 | 9493.48 | 10147.03 | 10163.94 | 7521.54 | 8318.77 | 8848.37 |
| Nucleus of the lateral olfactory tract' | 4338.75 | 4751.97 | 7541.17 | 4751.97 | 6198.22 | 5165.18 | 2995.81 | 5681.70 | 5888.31 | 5578.40 | 5578.40 | 3822.24 | 4855.27 |
| Cortical amygdalar area, anterior part' | 7289.93 | 6682.44 | 8059.42 | 5831.94 | 7370.93 | 6479.94 | 4292.96 | 8383.42 | 8221.42 | 8585.92 | 5426.95 | 6317.94 | 6925.43 |
| Cortical amygdalar area, posterior part' | 10248.91 | 10860.95 | 11817.96 | 10438.08 | 11651.04 | 11306.07 | 8802.27 | 10961.10 | 11996.01 | 12296.46 | 9592.35 | 10538.24 | 11940.37 |
| Piriform-amygdalar area' | 8318.71 | 9251.90 | 9651.84 | 8132.07 | 9705.16 | 9731.82 | 8931.95 | 9545.19 | 10718.34 | 10558.36 | 6825.61 | 8665.32 | 9785.15 |
| Postpiriform transition area' | 12218.95 | 12007.55 | 12916.57 | 12366.93 | 12134.39 | 13508.49 | 11753.87 | 12366.93 | 12388.07 | 12726.31 | 10781.42 | 12430.35 | 14502.07 |
| Field CA1' | 10774.30 | 10662.86 | 11829.77 | 9928.62 | 10810.36 | 11524.93 | 9102.60 | 10970.98 | 11000.48 | 11688.83 | 9548.39 | 10423.57 | 11987.11 |
| Field CA2' | 10628.42 | 9662.20 | 11456.61 | 9869.25 | 10076.30 | 10904.48 | 8903.03 | 12215.78 | 10283.34 | 9800.23 | 8419.92 | 7246.65 | 11939.72 |
| Field CA3' | 9858.78 | 10175.79 | 11660.42 | 8997.59 | 10233.90 | 10577.32 | 7703.17 | 10239.19 | 11147.93 | 11015.84 | 8765.12 | 10001.44 | 10778.09 |
| Dentate gyrus' | 10037.52 | 10143.34 | 12222.80 | 9915.82 | 10302.08 | 11005.82 | 7640.58 | 10937.03 | 9809.99 | 10741.25 | 9534.85 | 10354.99 | 11905.33 |
| Entorhinal area, lateral part' | 6884.85 | 5646.04 | 5751.96 | 5996.04 | 6175.64 | 6207.88 | 6014.46 | 6166.43 | 6051.30 | 5968.41 | 5747.36 | 5586.17 | 6553.28 |
| Entorhinal area, medial part, dorsal zone' | 9554.02 | 8450.92 | 9063.75 | 9131.16 | 8935.06 | 9566.27 | 8916.67 | 9265.99 | 8463.18 | 9302.76 | 9039.24 | 8873.77 | 10019.77 |
| Parasubiculum' | 13256.51 | 12268.23 | 12574.94 | 13426.90 | 13835.84 | 14790.04 | 12404.55 | 13017.96 | 10768.78 | 14347.02 | 12643.10 | 12540.86 | 12915.72 |
| Postsubiculum' | 12999.05 | 12503.17 | 13778.28 | 13424.09 | 13388.67 | 14451.26 | 12007.30 | 12821.95 | 13069.89 | 14380.42 | 12467.75 | 12821.95 | 14415.84 |
| Presubiculum' | 13790.56 | 13212.95 | 16101.03 | 12527.03 | 13682.26 | 15126.30 | 12851.94 | 13465.66 | 13357.35 | 15306.80 | 13140.75 | 13429.55 | 14368.18 |

| | | | | | | | | | | | | | |
|---|---|---|---|---|---|---|---|---|---|---|---|---|---|
| Subiculum' | 12684.48 | 12117.49 | 13656.48 | 12263.29 | 12538.69 | 13850.87 | 11550.50 | 12538.69 | 12311.89 | 13899.47 | 12311.89 | 12344.29 | 12830.28 |
| Cortical subplate' | 8446.90 | 8527.35 | 9734.05 | 7079.31 | 8688.24 | 9653.60 | 7320.65 | 9492.71 | 9412.26 | 9573.15 | 9090.47 | 6596.63 | 9010.03 |
| Claustrum' | 9893.82 | 9838.85 | 12367.27 | 9399.12 | 10828.23 | 9728.92 | 8464.71 | 9893.82 | 11432.85 | 11927.54 | 9179.26 | 7255.46 | 8189.88 |
| Endopirifor m nucleus' | 9255.86 | 8743.46 | 10237.04 | 8525.42 | 10269.75 | 10171.63 | 9005.11 | 10324.26 | 10651.32 | 11163.72 | 8143.84 | 8787.07 | 9462.99 |
| Lateral amygdalar nucleus' | 9205.91 | 10277.83 | 10593.10 | 10025.61 | 9931.03 | 10057.14 | 10530.05 | 10246.30 | 11507.39 | 13272.90 | 9426.60 | 9836.45 | 10719.21 |
| Basolateral amygdalar nucleus, anterior part' | 8146.43 | 8146.43 | 10819.47 | 8178.25 | 9705.70 | 9228.37 | 8019.14 | 9164.73 | 10469.43 | 9387.48 | 8337.36 | 7955.49 | 9260.19 |
| Basolateral amygdalar nucleus, posterior part' | 10993.48 | 9940.21 | 10368.10 | 7833.68 | 10598.51 | 11289.71 | 10763.08 | 11256.80 | 13659.56 | 12935.44 | 8393.23 | 8722.37 | 11289.71 |
| Basolateral amygdalar nucleus, ventral part' | 8856.46 | 9626.59 | 11744.44 | 8471.40 | 11616.09 | 10140.01 | 11359.38 | 10011.65 | 12193.68 | 12065.33 | 8471.40 | 8984.82 | 8920.64 |
| Basomedial amygdalar nucleus, anterior part' | 7033.45 | 7267.90 | 8322.92 | 7736.80 | 7814.95 | 8166.62 | 5861.21 | 7932.17 | 8166.62 | 8635.51 | 6017.51 | 6486.40 | 7932.17 |
| Basomedial amygdalar nucleus, posterior part' | 8904.53 | 11130.66 | 12066.28 | 10001.46 | 11130.66 | 11259.71 | 8936.79 | 11066.13 | 12195.33 | 12163.07 | 10162.77 | 10001.46 | 11453.29 |
| Posterior amygdalar nucleus' | 10363.87 | 8701.21 | 12248.21 | 8978.32 | 11610.86 | 11278.33 | 7537.36 | 10086.76 | 11278.33 | 12442.18 | 8867.48 | 9116.88 | 12109.65 |
| Caudoputa men' | 6637.72 | 5812.19 | 8237.85 | 6330.83 | 7746.01 | 7050.48 | 4831.21 | 7825.08 | 6868.22 | 6948.63 | 5737.14 | 6237.02 | 7082.65 |
| Nucleus accumbens' | 11257.00 | 9212.15 | 14293.29 | 10361.95 | 10031.47 | 10788.82 | 6382.41 | 12854.33 | 9611.48 | 9184.61 | 9336.08 | 10072.78 | 11973.04 |
| Olfactory tubercle' | 9830.38 | 7612.24 | 11099.08 | 8309.61 | 10141.25 | 9107.80 | 5755.39 | 10149.65 | 8864.14 | 8402.03 | 8133.17 | 8402.03 | 9813.57 |
| Lateral septal complex' | 10493.32 | 9297.39 | 14659.78 | 11361.33 | 9509.57 | 11390.27 | 7175.58 | 13994.31 | 10657.28 | 8593.33 | 10242.56 | 12402.95 | 15739.98 |
| Striatum-lik e amygdalar nuclei' | 4809.26 | 5771.11 | 7969.63 | 5816.91 | 6549.75 | 6320.74 | 3755.80 | 6824.57 | 5862.72 | 6229.14 | 5725.31 | 6045.93 | 7878.02 |
| Central amygdalar nucleus' | 7384.42 | 6836.46 | 8193.31 | 7593.17 | 7828.01 | 7880.19 | 5949.29 | 9628.45 | 9002.21 | 7932.38 | 5897.10 | 7332.23 | 7958.47 |
| Medial amygdalar nucleus' | 6188.12 | 5940.59 | 9947.40 | 6280.94 | 6234.53 | 6296.41 | 3357.05 | 7518.56 | 6590.35 | 7534.03 | 5878.71 | 7193.69 | 8493.19 |
| Globus pallidus, external segment' | 6052.58 | 5483.53 | 8225.30 | 6932.02 | 7216.54 | 6466.43 | 3931.59 | 7811.45 | 5897.39 | 7811.45 | 5612.86 | 6569.90 | 7992.51 |
| Globus pallidus, internal segment' | 5055.64 | 7058.82 | 10397.46 | 5914.15 | 6486.49 | 7631.16 | 2480.13 | 8489.67 | 5914.15 | 6486.49 | 8489.67 | 8489.67 | 10302.07 |
| Substantia innominata' | 8026.51 | 7428.67 | 10749.99 | 7860.45 | 7052.26 | 9011.83 | 4494.85 | 9997.16 | 7738.66 | 7971.16 | 7926.87 | 8347.57 | 10528.57 |
| Magnocellul ar nucleus' | 6638.88 | 6551.53 | 11618.04 | 8036.54 | 7250.36 | 9696.26 | 3494.15 | 10307.74 | 6900.94 | 6289.47 | 9172.14 | 9521.55 | 9870.97 |

| | | | | | | | | | | | | | |
|---|---|---|---|---|---|---|---|---|---|---|---|---|---|
| Medial septal nucleus' | 14986.11 | 15913.08 | 16994.55 | 20934.20 | 14136.38 | 23251.64 | 6643.33 | 23792.38 | 14522.62 | 12436.92 | 8265.53 | 19003.00 | 13132.16 |
| Diagonal band nucleus' | 12389.65 | 9752.50 | 13683.35 | 12439.41 | 12091.11 | 13235.53 | 5970.92 | 15225.84 | 12389.65 | 9205.16 | 9105.65 | 10250.08 | 11941.84 |
| Triangular nucleus of septum' | 10514.49 | 11111.90 | 18997.77 | 12306.73 | 9917.08 | 12545.70 | 6452.07 | 17444.49 | 8961.21 | 8363.80 | 11350.87 | 14815.87 | 15413.29 |
| Bed nuclei of the stria terminalis' | 11753.89 | 6309.99 | 7943.16 | 5518.14 | 6804.89 | 7720.45 | 4874.77 | 7349.28 | 9205.15 | 7027.59 | 5419.16 | 7349.28 | 7126.57 |
| Ventral group of the dorsal thalamus' | 7220.74 | 7085.40 | 9959.36 | 6894.33 | 6098.22 | 8279.57 | 3725.80 | 8757.23 | 6233.56 | 7690.44 | 6902.29 | 8104.42 | 9959.36 |
| Subparafascicular nucleus' | 6778.20 | 9037.60 | 12991.55 | 6401.63 | 6401.63 | 9225.88 | 5648.50 | 5271.93 | 7343.05 | 8661.03 | 9602.45 | 11108.72 | 12238.42 |
| Subparafascicular area' | 11001.80 | 8975.15 | 13317.97 | 8975.15 | 10133.24 | 11870.36 | 6079.94 | 11001.80 | 8396.11 | 9264.67 | 15634.14 | 15344.62 | 9554.20 |
| Peripeduncular nucleus' | 12508.89 | 14295.87 | 15487.20 | 11317.57 | 14295.87 | 10721.91 | 11913.23 | 12508.89 | 10721.91 | 13700.21 | 7147.94 | 11317.57 | 10721.91 |
| Geniculate group, dorsal thalamus' | 11155.38 | 10885.93 | 13957.70 | 10508.69 | 12556.54 | 14281.04 | 8945.86 | 11586.51 | 12664.32 | 10993.71 | 9484.77 | 12340.98 | 9808.11 |
| Lateral group of the dorsal thalamus' | 7040.13 | 8077.50 | 11003.12 | 7378.15 | 7261.59 | 9382.95 | 4301.01 | 9452.89 | 6748.73 | 7844.38 | 6923.57 | 7925.97 | 10991.46 |
| Anterior group of the dorsal thalamus' | 8987.17 | 7358.83 | 10302.37 | 7233.58 | 7812.89 | 9378.60 | 3742.05 | 8987.17 | 7061.35 | 7108.32 | 7233.58 | 8783.63 | 8799.29 |
| Medial group of the dorsal thalamus' | 11516.31 | 11441.77 | 12541.22 | 10901.36 | 9969.62 | 11702.66 | 5720.88 | 12858.01 | 10137.33 | 10211.87 | 9522.38 | 12727.57 | 15709.14 |
| Midline group of the dorsal thalamus' | 10925.79 | 12425.41 | 14567.72 | 12517.22 | 9273.15 | 13129.31 | 5845.45 | 15149.21 | 9671.01 | 7987.76 | 11231.84 | 13527.17 | 16342.78 |
| Intralaminar nuclei of the dorsal thalamus' | 10061.56 | 11081.14 | 12556.83 | 10410.37 | 9739.59 | 11751.91 | 5688.14 | 12503.17 | 8344.39 | 9551.78 | 8129.74 | 11322.61 | 15213.08 |
| Reticular nucleus of the thalamus' | 7318.36 | 7237.35 | 9262.72 | 7723.44 | 6211.16 | 8560.59 | 4023.75 | 8236.53 | 7183.34 | 7912.47 | 7696.43 | 7777.45 | 8749.63 |
| Intergeniculate leaflet of the lateral geniculate complex' | 10534.59 | 11705.10 | 10534.59 | 6437.81 | 16972.40 | 9364.08 | 8193.57 | 9364.08 | 11705.10 | 14046.12 | 9949.34 | 7608.32 | 9364.08 |
| Ventral part of the lateral geniculate complex' | 10465.58 | 9967.22 | 12957.39 | 9468.86 | 11661.65 | 10066.89 | 7276.07 | 10465.58 | 13455.75 | 11860.99 | 9070.17 | 9269.52 | 10864.27 |
| Subgeniculate nucleus' | 1645.52 | 6582.07 | 14809.66 | 9873.11 | 14809.66 | 14809.66 | 3291.04 | 18100.70 | 8227.59 | 14809.66 | 3291.04 | 4936.55 | 4936.55 |
| Medial habenula' | 11474.68 | 14089.16 | 13072.42 | 8714.94 | 9876.94 | 13362.91 | 7262.45 | 11038.93 | 9876.94 | 8569.69 | 10312.68 | 16558.39 | 18446.63 |

| | | | | | | | | | | | | | |
|---|---|---|---|---|---|---|---|---|---|---|---|---|---|
| Lateral habenula' | 12138.28 | 12579.67 | 13352.11 | 11696.89 | 11586.54 | 12579.67 | 7393.32 | 13683.15 | 9931.32 | 10593.41 | 9379.58 | 12690.02 | 17766.03 |
| Periventricular zone' | 12101.92 | 10968.63 | 15178.00 | 13356.64 | 9592.49 | 13963.76 | 6192.62 | 15420.85 | 9471.07 | 9228.22 | 11413.85 | 12709.04 | 14206.61 |
| Periventricular region' | 9783.14 | 10633.18 | 13322.38 | 10540.45 | 9814.05 | 11823.23 | 5857.52 | 13322.38 | 9659.50 | 8794.01 | 9134.02 | 12704.18 | 13507.85 |
| Anterior hypothalamic nucleus' | 8866.36 | 7549.58 | 11104.90 | 8646.90 | 7681.25 | 8954.15 | 3204.18 | 9480.86 | 8515.22 | 7374.00 | 7330.11 | 8515.22 | 10973.22 |
| Mammillary body' | 12198.38 | 11548.76 | 15121.66 | 11296.13 | 10069.07 | 13461.52 | 7687.14 | 13858.51 | 8986.38 | 11909.66 | 9960.80 | 13750.24 | 12667.54 |
| Medial preoptic nucleus' | 8212.07 | 7596.16 | 12318.11 | 9922.92 | 6911.83 | 7596.16 | 4585.07 | 10196.65 | 8554.24 | 5474.71 | 8691.11 | 11496.90 | 17450.65 |
| Dorsal premammillary nucleus' | 9984.37 | 13214.60 | 11746.31 | 8516.08 | 9690.71 | 10278.03 | 7928.76 | 7928.76 | 10278.03 | 6460.47 | 9397.05 | 11452.66 | 12920.95 |
| Ventral premammillary nucleus' | 9210.55 | 10315.82 | 11421.08 | 10500.03 | 10315.82 | 10500.03 | 7736.86 | 9394.76 | 8473.71 | 11052.66 | 8473.71 | 11605.29 | 15105.30 |
| Paraventricular hypothalamic nucleus, descending division' | 12802.72 | 6668.08 | 13336.16 | 7201.53 | 8535.15 | 10402.21 | 4801.02 | 9602.04 | 4801.02 | 8535.15 | 9868.76 | 15203.23 | 12535.99 |
| Ventromedial hypothalamic nucleus' | 11845.14 | 11119.93 | 15108.60 | 12026.45 | 11361.67 | 12026.45 | 5801.70 | 13476.87 | 10032.11 | 11724.27 | 10152.98 | 15652.51 | 15833.81 |
| Posterior hypothalamic nucleus' | 11445.63 | 11047.02 | 13438.65 | 8826.23 | 10022.04 | 13780.31 | 8199.85 | 12869.21 | 9395.66 | 10249.81 | 9110.95 | 11502.57 | 14235.85 |
| Hypothalamic lateral zone' | 8310.76 | 7925.26 | 9682.44 | 7862.50 | 6526.68 | 9368.66 | 3612.98 | 9296.93 | 6867.36 | 7772.85 | 7037.70 | 8463.17 | 10570.00 |
| Zona incerta' | 7495.29 | 7151.66 | 9943.61 | 6357.04 | 6464.42 | 7946.29 | 3285.90 | 8268.44 | 6936.90 | 7430.86 | 7259.05 | 7989.25 | 9148.98 |
| Median eminence' | 12830.13 | 12830.13 | 10157.19 | 11760.95 | 8018.83 | 12830.13 | 8018.83 | 12295.54 | 7484.24 | 10691.77 | 14433.89 | 14968.48 | 12295.54 |
| Superior colliculus, sensory related' | 13192.88 | 11479.05 | 13674.89 | 12300.26 | 12211.00 | 13960.53 | 10050.87 | 13139.32 | 11050.60 | 14014.08 | 11086.30 | 12228.85 | 12710.86 |
| Inferior colliculus' | 11350.75 | 11166.76 | 12482.99 | 11633.81 | 11796.57 | 12801.43 | 11463.97 | 12107.94 | 11159.68 | 11902.72 | 11046.46 | 11860.26 | 12801.43 |
| Substantia nigra, reticular part' | 10178.15 | 9624.24 | 12647.68 | 9324.20 | 10178.15 | 10016.59 | 7639.38 | 9601.16 | 10062.75 | 10270.47 | 8862.61 | 10016.59 | 10755.14 |
| Ventral tegmental area' | 10196.46 | 8676.49 | 13173.06 | 9436.47 | 6966.52 | 10196.46 | 7093.19 | 10893.11 | 8676.49 | 10513.12 | 8993.15 | 10766.44 | 12223.08 |
| Superior colliculus, motor related' | 11918.18 | 10826.82 | 13804.56 | 10862.96 | 11744.72 | 13060.13 | 9439.14 | 11491.76 | 11549.58 | 12597.57 | 10364.26 | 11527.89 | 12221.74 |
| Periaqueductal gray' | 13603.70 | 12811.00 | 13671.89 | 13586.65 | 13970.21 | 15172.04 | 9947.06 | 14294.11 | 11881.93 | 14413.44 | 12112.06 | 13884.98 | 14396.39 |
| Precommissural nucleus' | 10591.83 | 12945.57 | 13416.32 | 7061.22 | 8002.72 | 12004.08 | 7531.97 | 10356.46 | 10591.83 | 11062.58 | 8002.72 | 12710.20 | 15299.31 |
| Anterior pretectal nucleus' | 8924.21 | 8381.52 | 11456.75 | 6904.20 | 6602.71 | 8411.67 | 4763.60 | 7145.40 | 6843.90 | 8713.16 | 8321.22 | 8411.67 | 9255.85 |
| Medial pretectal area' | 11653.62 | 8324.02 | 10821.22 | 11653.62 | 5826.81 | 14983.23 | 5826.81 | 10821.22 | 9156.42 | 9156.42 | 9156.42 | 11653.62 | 9988.82 |

| | | | | | | | | | | | | | |
|---|---|---|---|---|---|---|---|---|---|---|---|---|---|
| Substantia nigra, compact part' | 9662.49 | 8309.74 | 13140.98 | 6183.99 | 7246.86 | 9082.74 | 5700.87 | 10338.86 | 8986.11 | 9179.36 | 7150.24 | 8599.61 | 9469.24 |
| Pedunculopontine nucleus' | 12689.73 | 11595.78 | 14658.82 | 12821.00 | 13827.43 | 15577.73 | 13083.54 | 14790.09 | 14046.21 | 14615.06 | 13302.33 | 14790.09 | 14046.21 |
| Interfascicular nucleus raphe' | 24062.99 | 18047.24 | 21749.24 | 19898.24 | 16658.99 | 19435.49 | 12031.49 | 15733.49 | 11568.74 | 23600.24 | 15270.74 | 22674.74 | 17584.49 |
| Interpeduncular nucleus' | 23017.46 | 23017.46 | 17263.10 | 17263.10 | 23017.46 | 25894.65 | 5754.37 | 14385.91 | 14385.91 | 25894.65 | 23017.46 | 17263.10 | 8631.55 |
| Rostral linear nucleus raphe' | 25873.98 | 28597.56 | 20426.83 | 17022.36 | 14298.78 | 29959.35 | 21107.72 | 26554.88 | 10894.31 | 35406.50 | 12936.99 | 24512.20 | 14979.67 |
| Central linear nucleus raphe' | 15867.77 | 22314.05 | 20330.58 | 14876.03 | 22809.92 | 23305.79 | 18842.98 | 21322.31 | 12892.56 | 25785.12 | 23305.79 | 16859.50 | 15867.77 |
| Dorsal nucleus raphe' | 22130.47 | 8694.11 | 16334.40 | 10538.32 | 16597.85 | 9484.49 | 17124.77 | 13963.27 | 21867.01 | 26609.26 | 20549.72 | 11592.15 | 7376.82 |
